# The role of binocular disparity and attention in the neural representation of multiple moving stimuli in the visual cortex

**DOI:** 10.1101/2023.06.25.546480

**Authors:** Anjani Sreeprada Chakrala, Jianbo Xiao, Xin Huang

**Author notes:** Correspondence should be addressed to: Xin Huang, Department of Neuroscience, University of Wisconsin, Madison, WI 53705, USA. These authors contributed equally to this work (Anjani Sreeprada Chakrala, Jianbo Xiao).

## Abstract

Segmenting visual scenes into distinct objects and surfaces is a fundamental visual process, with stereoscopic depth and motion serving as crucial cues. However, how the visual system uses these cues to segment multiple objects is not fully understood. We investigated how neurons in the middle-temporal (MT) cortex of macaque monkeys represent overlapping surfaces at different depths, moving in different directions. Neuronal activity was recorded from three male monkeys during discrimination tasks under varying attention conditions. We found that neuronal responses to overlapping surfaces showed a robust bias toward the binocular disparity of one surface over the other. The disparity bias of a neuron was positively correlated with the neuron’s disparity preference for a single surface. In two animals, neurons preferring near disparities of single surfaces (near neurons) showed a near bias for overlapping stimuli, while neurons preferring far disparities (far neurons) showed a far bias. In the third animal, both near and far neurons displayed a near bias, though the near neurons showed a stronger near bias. All three animals exhibited an initial near bias across neurons relative to the average of the responses to the individual surfaces. Although attention modulated neuronal responses, the disparity bias was not caused by attention. We also found that the effect of attention was consistent with object-based, rather than feature-based attention. We proposed a model in which the pool size of the neuron population that weighs the responses to individual stimulus components can be variable. This model is a novel extension of the standard normalization model and provides a unified explanation for the disparity bias across animals. Our results reveal how MT neurons encode multiple stimuli moving at different depths and present new evidence of response modulation by object-based attention. The disparity bias allows subgroups of neurons to preferentially represent individual surfaces of multiple stimuli at different depths, thereby facilitating segmentation.

## Introduction

Natural scenes often contain multiple entities within a three-dimensional (3D) space. Segmenting these visual scenes into distinct objects and surfaces is a fundamental process of vision. Among various visual features, stereoscopic depth and motion cues are particularly important for distinguishing different objects from one another and segregating figural objects from their background (Gibson, 1950; Wagemans et al., 2012). While the visual system can rely on depth or motion cues alone for segmentation, combining these cues further enhances the process (Snowden and Rossiter, 1999; Goutcher 2016). For example, it is easier to segment overlapping surfaces moving in different directions when they are at different depths rather than at the same depth (Qian et al., 1994; Hibbard and Bradshaw, 1999; Greenwood and Edwards, 2006). To understand the neural basis of natural vision, it is important to determine how the visual system represents multiple visual stimuli that differ in various features and how depth cues facilitate the segmentation of multiple stimuli.

The middle-temporal cortex (MT) of primates is crucial for processing motion and depth information (Born and Bradley, 2005; Britten, 2003; Pasternak and Tadin, 2020). Most MT neurons are selective to motion direction and binocular disparity (Maunsell and van Essen, 1983a, 1983b; Albright, 1984; DeAngelis et al., 1998; DeAngelis and Newsome, 1999). Although the representation of binocular disparity for a single surface by MT neurons is well understood (Cumming and Parker, 1999; DeAngelis and Uka, 2003; Uka and DeAngelis, 2003, 2004; Cumming and DeAngelis, 2001), the understanding of how MT neurons represent multiple stimuli at different disparities remains incomplete.

One previous study showed that the responses of MT neurons to two overlapping stimuli moving in opposite (the preferred and null) directions became stronger as the disparity difference between the stimuli increased, suggesting reduced opponent inhibition between motion signals with increased disparity difference (Bradley et al., 1995). Other studies showed that MT neurons can represent structure-from-motion (Bradley et al., 1998; Dodd et al., 2001; Krug et al., 2013) and relative disparity (Krug and Parker, 2011) using overlapping surfaces moving in opposite directions at different disparities. Uka and DeAngelis (2006) also investigated MT neurons’ selectivity for relative disparity using center-surround stimuli moving in the same (preferred) directions of the neurons at different disparities. However, to gain a general understanding of how MT neurons encode multiple stimuli at different disparities, it is essential to examine visual stimuli across different motion directions and with varying angular separations between them. We address this question by measuring the direction tuning curves of MT neurons to two overlapping surfaces located at different disparities – one near and one far from the fixation plane – moving in various directions across 360° with different angular separations.

We anticipate that MT neurons will follow one of three encoding rules for multiple stimuli at different disparities. First, they may average the responses to individual stimulus components at different disparities, treating both components equally. Second, they may show a consistent bias toward one stimulus disparity (e.g. the near surface), regardless of individual neurons’ disparity preferences for single surfaces, which could be beneficial for efficiently representing disparity information or performing tasks like figure-ground segregation in natural environments (Sprague et al., 2015; Manning et al., 2024). Third, MT neurons may bias their response toward the disparity component closer to their preferred disparity, enhancing the extraction of information from the more preferred disparity component. The primary goal of this study is to determine the encoding rules for multiple disparities of moving stimuli.

Attention can bias neuronal responses elicited by multiple two-dimensional (2D) stimuli in the receptive field (RF), in favor of the attended stimulus (Desimone and Duncan, 1995; Reynolds et al., 1999; Treue and Maunsell, 1996; Ferrera and Lisberger, 1997; Recanzone and Wurtz, 2000). MT neuron responses can be influenced by feature-based attention, such as attention to motion direction (Treue and Martinez-Trujillo, 1999; Martinez-Trujillo and Treue, 2004) and binocular disparity (Ruff and Born, 2015). Attention can also select a specific visual object (Duncan, 1984; Cavanagh et al., 2023; Valdes-Sosa et al., 1998, 2000; Mitchell et al., 2003, 2004). Wannig et al. (2007) demonstrated that MT neuron responses could be modulated by object-based attention directed at a moving surface composed of two overlapping 2D stimuli. However, the effects of selective attention on the neural representation of multiple stimuli located at different depths remain underexplored. When subjects are instructed to attend to the disparity of one of two overlapping moving surfaces at different depths, it is unclear whether the effect of attention is feature-based and restricted to disparity or if it extends to all the features of the entire surface, indicating object-based attention. This study aims to address this question as its secondary goal.

We recorded neuronal activities in MT of three macaque monkeys while they performed discrimination tasks under different attention conditions. We found that neuronal responses to overlapping surfaces at different depths showed a robust bias toward the disparity of one of the surfaces. Across all animals, a neuron’s disparity bias in response to two surfaces was positively correlated with its disparity preference for a single surface. Notably, the combined responses of near and far neurons of all three animals showed an initial bias toward the near surface. Additionally, we found that the disparity bias was not caused by attention, although attention modulated neuronal responses in a manner consistent with object-based attention. We proposed a model that extended the standard normalization model to account for our findings.

## Results

We asked how the primate visual system utilizes disparity and motion cues to segment multiple visual stimuli. To address this question, we characterized how neurons in area MT of macaque monkeys represent overlapping stimuli located at different depths and moving simultaneously in different directions, while monkeys performed discrimination tasks under different attention conditions.

### Behavioral task and performance

We trained two monkeys (B and G) to perform a direction discrimination task. The visual stimuli were two overlapping random-dot patches moving in two directions separated by either 60° or 120°. By randomly interleaving trials with 60° and 120° direction separations, monkeys are prevented from performing the task by merely identifying the direction of one component and guessing the other based on a fixed separation. The two direction separations, 120° and 60°, also offered relatively easier and more challenging scenarios for segmentation, respectively. One patch was located at a near disparity (−0.1°) and the other patch at a far disparity (0.1°). The monkey’s task was to report the motion direction of the patch at a cued depth (either the near or far disparity) by making a saccadic eye movement to one of twelve reporting targets (12 AFC task) (Fig. 1A, B). We varied the vector average (VA) direction of the overlapping stimuli across trials. The cue-near trials (Fig. 1A) and cue-far trials (Fig. 1B) were randomly interleaved. To perform the task well, the monkeys need to segment two overlapping surfaces and attend to the cued surface to discriminate its motion direction. We therefore refer to the cue-near and cue-far trials as “Attend Near” and “Attend Far” trials, respectively. We also randomly interleaved trials in which only one of the two patches was presented at either the near (−0.1°) or far (0.1°) disparity.

**Figure 1.**
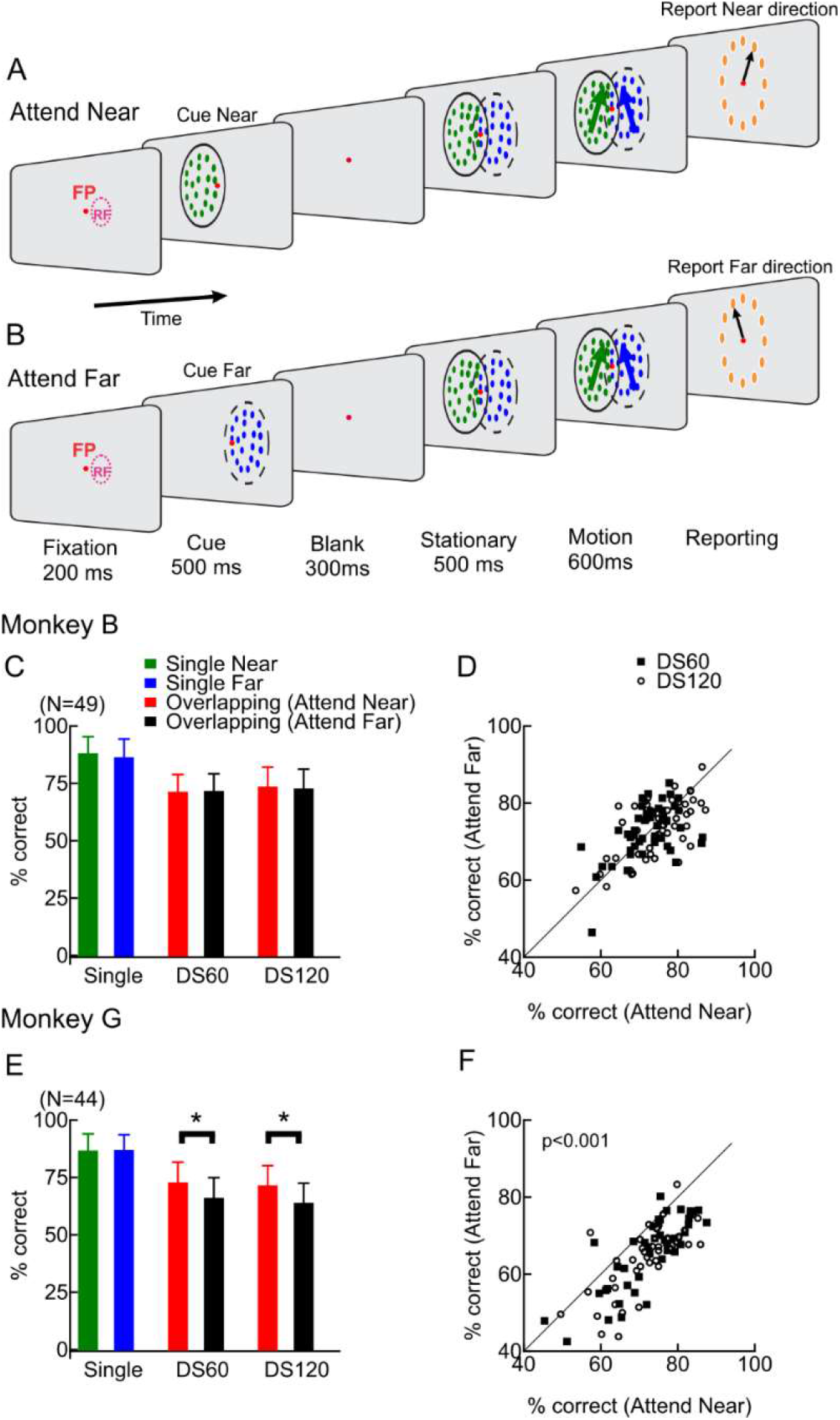
The behavioral task and performance. **A, B**. Visual stimuli were overlapping random-dot patches located at different depths and moving in different directions. Each trial started with fixation on a spot at the center of the monitor. Then, as a cue, a stationary random-dot patch (RDP) was presented for 500 ms at either the near (−0.1°) (A) or the far (0.1°) disparity (B) for 500 ms. After a 300-ms blank period, overlapping random-dot stimuli were turned on, initially stationary for 500 ms, and then moved in two directions for 600 ms. The direction separation (DS) between the two motion directions was either 60° or 120°. After the visual stimuli were turned off, 12 reporting targets were illuminated. To receive a juice reward, the monkey needed to make a saccadic eye movement to one of the targets in the direction that matched the motion direction of the cued surface of overlapping stimuli. **A**. Cue-near (Attend-Near) trial. **B**. Cue-far (Attend-Far) trial. **C. D**. Behavioral performance of monkey B. C shows the mean correct percentages across sessions for single and overlapping stimuli. D compares the performance between Attend Near and Attend Far conditions. Each dot in D represents the trial-averaged correct percentage in one session. **E. F**. Behavioral performance of monkey G, following the same convention as in C and D. Error bars in C and E represent standard deviation. (*) in E indicates p < 0.001 (paired signed-rank test).

We recorded neuronal activities in area MT while the monkeys performed the task (49 experimental sessions for monkey B, 44 sessions for monkey G). Both monkeys performed the task well. The mean correct rate across sessions was 75% (sd = 6.6) for monkey B (single-surface 87%, two-surface 72%), and 72% (sd = 8.1) for monkey G (single-surface 87%, two-surface 68%). As expected, both monkeys performed significantly better in the single-surface trials than in two-surface trials (signed rank test, p < 10^−8^) (Fig. 1C, E). Notably, while monkey B performed equally well in the “Attend Near” and “Attend Far” trials (signed rank test, p = 0.56) (Fig. 1C, D), monkey G performed significantly better at discriminating the direction of the near-surface (mean = 72% correct) than the far-surface (mean = 65% correct) of the overlapping stimuli (one-sided signed rank test, p= 4.3×10^−13^) (Fig. 1E, F).

### Direction tuning of example MT neurons to overlapping stimuli moving at different depths

Figure 2 shows the direction tuning curves of four example MT neurons to overlapping stimuli moving in different directions at different depths. The red and black curves indicate the responses under attend-near and attend-far conditions, respectively. The overlapping stimuli had four configurations (Fig. 2A-D). The motion direction at the near surface could either be at the clockwise (A, C) or counter-clockwise side (B, D) of the two motion directions, and the direction separation between the two directions was either 60° (A, B) or 120° (C, D). We referred to the overlapping stimuli as the bi-directional stimuli. Figure 2 also shows the direction tuning curves to single motion directions of the constituent near-surface (green) and far-surface (blue) when they were presented individually. We classified a neuron as a near-preferred (or far-preferred) neuron if the maximum response of the neuron’s direction tuning curve to the near (or far) component was greater than that to the other component.

**Figure 2.**
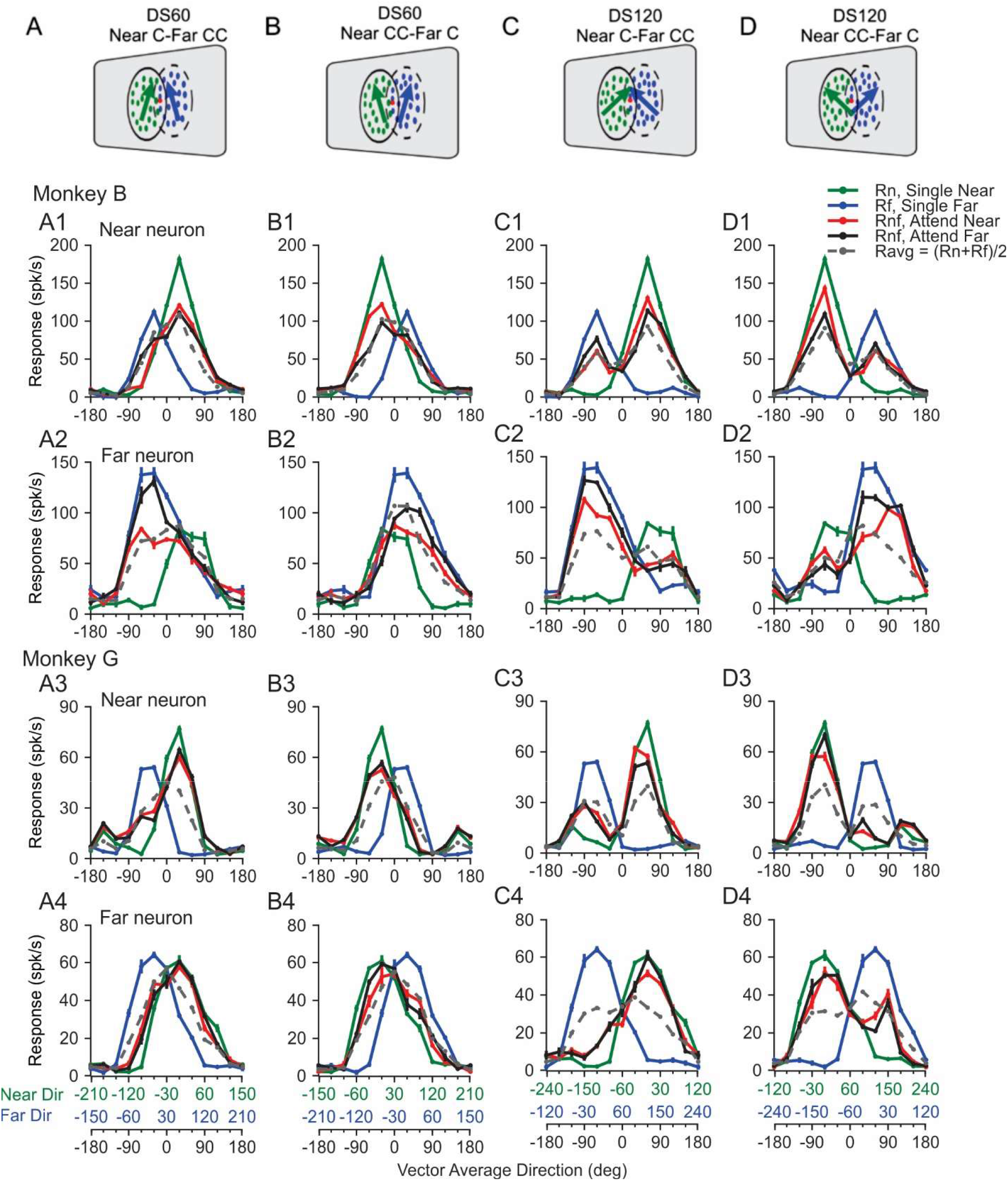
Direction tuning curves of example MT neurons in response to overlapping bi-directional stimuli and the constituent stimulus components. **A-D. Stimlus configurations.** The Direction separation between the two stimulus components was either 60° (A, B) or 120° (C, D). The motion direction at the near surface can either be at the clockwise (A, C) or counter-clockwise side (B, D) of the two motion directions. Responses of a near neuron **(A1-D1)** and a far neuron **(A2-D2)** from monkey B. Responses of a near neuron **(A3-D3)** and a far neuron **(A4-D4)** from monkey G. The red and black curves indicate the responses under attend-near and attend-far conditions, respectively. The green and blue curves indicate the responses to single motion directions of the near-surface and far-surface, respectively, when they were presented individually. The gray dash line indicates the average of the responses to the component directions (i.e. average of the green and blue curves). The abscissas in green and blue show the directions for the near component and the far component, respectively, of the corresponding bi-directional stimuli for which the vector average (VA) direction is shown by the black abscissa. The green and blue axes are shifted by 60° relative to each other in columns A and B, and by 120° in columns C and D. VA direction of 0° is aligned with each neuron’s preferred direction (PD). Error bars indicate the standard error.

Figure 2A1-D1 shows the responses of a near-preferred neuron, referred to as “near neuron”, from monkey B. The response tuning curve to the bi-directional stimuli under the attend-near condition (red curve) was biased toward the near component (green curve). By definition, the near neuron responded more strongly to the near component than the far component, and therefore the average of the component responses (gray) had a peak toward the near component. However, the red curve showed more bias toward the near component than expected by the component average. This near bias was found at both direction separations of 60° and 120°, regardless of whether the near direction was at the clockwise or counterclockwise side of the two motion directions. Under the attend-far condition, the response tuning to the bi-directional stimuli still showed a near bias in most of the stimulus configurations (black curve, Fig. 2A1, C1, D1), but to a lesser degree than that under the attend-near condition.

Figure 2A2-D2 shows the responses of a far neuron from monkey B. The response tuning curve to the bi-directional stimuli under the attend-far condition (black curve) was biased toward the far component (blue curve). Under the attend-near condition, the response tuning to the bidirectional stimuli still showed a far bias in some stimulus configurations, albeit weaker (red curve, Fig. 2B2, C2, D2).

Figures 2A3-D3 and 2A4-D4 show the responses of an example near neuron and a far neuron from monkey G. The near neuron showed a strong near bias under either attend-near or attend-far conditions (Fig. 2A3-D3). Remarkably, the far neuron also showed a near bias regardless of the attention (Fig. 2A4-D4). The responses to the bi-directional stimuli of these two neurons showed small differences between the attend-near and attend-far conditions.

We previously found that some MT neurons showed response bias toward one direction component of two overlapping stimuli moving in different directions within a 2-dimensional plane (Xiao and Huang, 2015). We noticed that some MT neurons also showed this directional bias to stimuli moving at different depths. For example, the far neuron from monkey B showed a stronger far bias when the far direction was at the counter-clockwise side of the two motion directions (the black curve was very close to the blue curve, Fig. 2A2, C2) than when the far direction was at the clockwise side (Fig. 2B2, D2). Our experimental design allowed us to separate the effect of disparity bias from the directional bias as shown below.

### Population-averaged direction tuning curves

Figure 3 shows the tuning curves to the bi-directional stimuli averaged across near neurons and far neurons for each animal. The direction tuning curve of each neuron was first fitted with a spline, normalized, and then shifted circularly such that the neuron’s preferred direction (PD) was aligned with the VA direction 0° of the bi-directional stimuli. To examine disparity bias and balance potential directional bias, we averaged the response tuning curves to the bi-directional stimuli when the near component was at the clockwise side (Near C/Far CC) (e.g. Fig. 2A1) and at the counter-clockwise (Near CC/Far C) side (e.g. Fig. 2B1) of the two component directions. Before averaging, we flipped the tuning curve of the stimulus configuration when the near component was at the counter-clockwise side (Near CC/Far C) relative to VA 0°, so in the flipped tuning curve, the near component was also at the clockwise side of two component directions. Averaging these two tuning curves preserved the disparity bias but cancelled out the directional bias. As a result, a response bias toward positive VA values indicates a bias toward the near component. The tuning curves of each neuron were then averaged across neurons for 60° (Fig. 3A, B) and 120° (Fig. 3C, D) direction separations.

**Figure 3.**
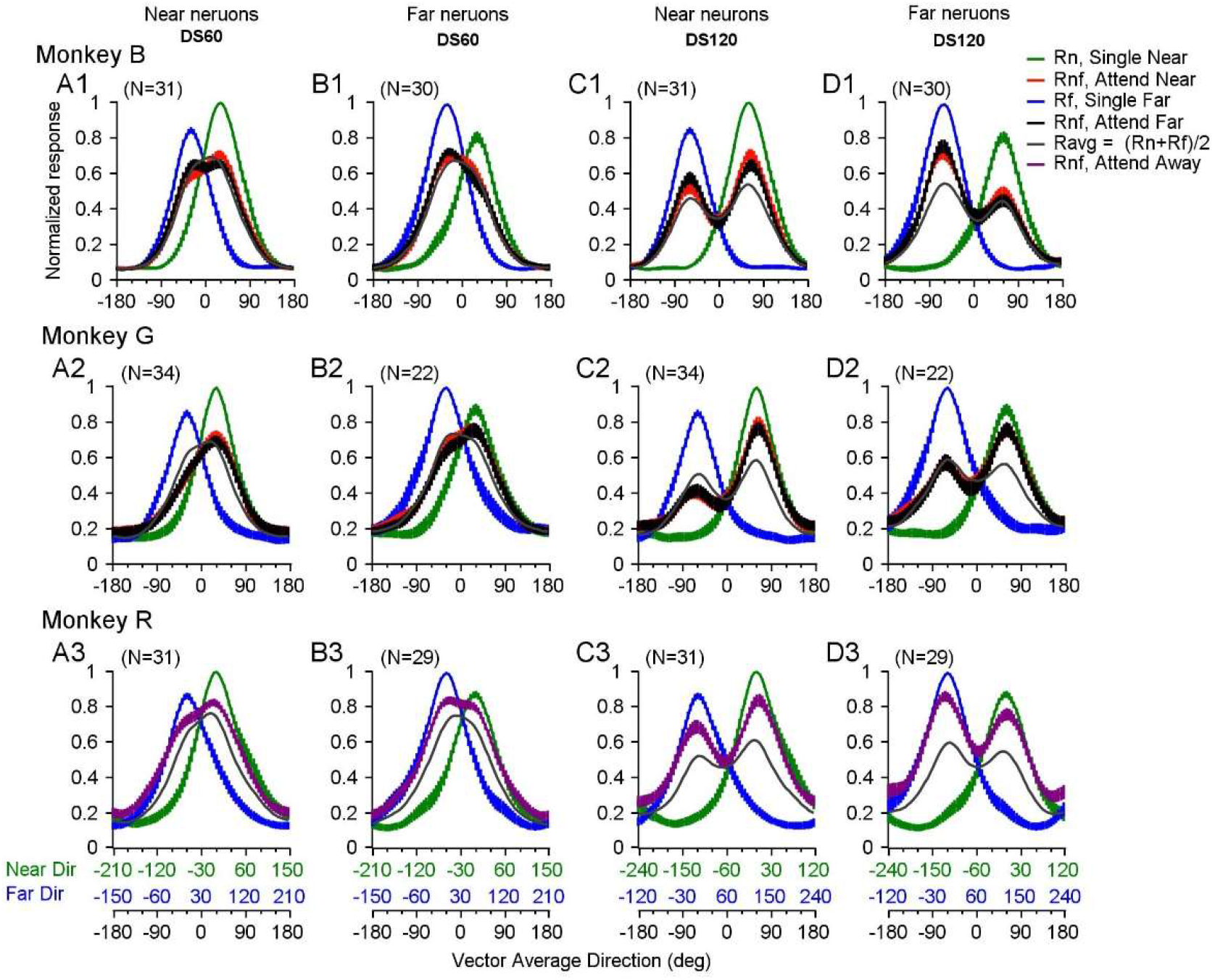
Population-averaged direction tuning curves to overlapping stimuli moving at different depths. Tuning curves to the bi-directional stimuli with 60° direction separation (DS60) (**A, B**) and 120° direction separation (DS120) (**C, D**), and the corresponding component directions. **A, C**. Tuning curves averaged across near neurons. **B, D**. Tuning curves averaged across far neurons. **A1-D1**. Responses from monkey B. **A2-D2**. Responses from monkey G. **A3-D3**. Responses from monkey R when attention was directed away from the RFs. The width of the curve for *R*_*n*_, *R*_*f*_, and *R*_*nf*_ represents the standard error. The tuning curve of each neuron was spline-fitted, normalized, and shifted circularly such that each neuron’s PD was aligned with VA 0°. The tuning curves to the bi-directional stimuli that had the near component at the clockwise side (Near C/Far CC) and at the counter-clockwise side (Near CC/Far C) of the two component directions were averaged. The tuning curves were then averaged across neurons. As a result, in all panels, the near component is always at the clockwise side of the two motion directions.

For monkey B, the population-averaged tuning curves to the bi-directional stimuli showed a bias toward neurons’ preferred disparity – the near neurons showed a bias toward the near component (Fig. 3A1, C1), and the far neurons showed a bias toward the far component (Fig. 3B1, D1). The disparity bias was modulated by attention – the near neurons showed a stronger near bias when attention was directed to the near surface (Fig. 3A1, C1) and the far neurons showed a stronger far bias when attention was directed to the far surface (Fig. 3B1, D1). However, the disparity bias cannot be explained by attention modulation. For example, far neurons still showed a clear far bias even when attention was directed to the near surface (red curve in Fig. 3D1).

In contrast, for monkey G, the population-averaged tuning curves to the bi-directional stimuli showed a robust bias toward the near component for both the near neurons (Fig. 3A2, C2) and the far neurons (Fig. 3B2, D2). Even in trials when attention was directed to the far surface, the far neurons still showed a bias toward the near component (black curves in Fig. 3B2, D2). In comparison to the attention modulation found in the data from monkey B, the effect of attention modulation found from monkey G was small.

Given the apparent discrepancy in the response tuning curves and the magnitude of attention modulation between monkeys B and G, we recorded from area MT of a third monkey R. We asked whether MT neurons showed disparity biases without the involvement of attention. We trained monkey R to perform a demanding fine direction-discrimination task in the visual hemifield opposite from the RFs of the recorded MT neurons, while we presented the bi-directional stimuli in the RFs. The results from monkey R showed a similar disparity bias as found in monkey B – the near neurons tended to show a near bias (Fig. 3A3, C3) and the far neurons tended to show a far bias (Fig. 3B3, D3).

### Quantifying the disparity bias and the effect of attention

To quantify the response bias toward either the near or far component, we fitted the tuning curve to the bi-directional stimuli of each neuron (the red and black curves illustrated in Fig. 2) as a function of the neuron’s responses to the individual stimulus components (the green and blue curves illustrated in Fig. 2). We found that a model of weighted summation plus a nonlinear interaction term, referred to as the SNL model (Eq. 1), provided a good fit of the MT responses. The percentage of variances (PV) (Eq. 13 in Methods) accounted for by the SNL model were 93%, 83%, and 88% for monkeys B, G, and R, respectively. The SNL model fit was significantly better than a linear weighted summation model replacing the nonlinear interaction term with a constant (one-sided signed-rank test, p < 10^−7^).

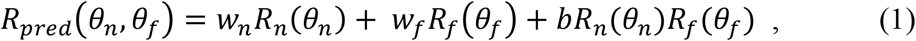

in which *R*_*pred*_ is the model-predicted response to bi-directional stimuli; *θ*_*n*_ and *θ*_*f*_ are the motion directions of the near and far components; *R*_*n*_ and *R*_*f*_ are the responses to single motion directions at the near and far surfaces; *w*_*n*_ and *w*_*f*_ are the response weights for the near and far components, respectively; *b* is a coefficient for the multiplicative interaction between the two component responses; *w*_*n*_, *w*_*f*_, and *b* are model parameters.

Using the fitted response weights for the near and far components, we defined a weight bias index (*WBI*) as:

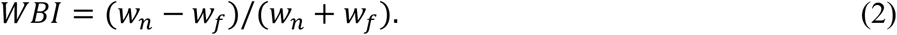

A positive *WBI* indicates a bias toward the near component, whereas a negative *WBI* indicates a bias toward the far component. Figure 4 shows the *WBI* of the near and far neurons. In the scatter plots (Fig. 4A, B), each neuron contributed to four points, from direction separations of 60° and 120°, and stimulus configurations of Near C/Far CC and Near CC/Far C. For monkey B, near neurons had significantly larger *WBI* than the far neurons, regardless of whether attention was directed to the near or far surface (one-sided signed-rank test, p < 0.001) (Fig. 4A). When attention was directed to the near surface, near neurons showed a significant near bias (mean *WBI* = 0.12, p = 1.43×10^−8^, one-sided signed-rank test), and far neurons showed a significant far bias (mean *WBI* = -0.17, p = 1.95×10^−8^). When attention was directed to the far surface, far neurons showed a significant far bias (mean *WBI* = -0.30, p= 5.74×10^−15^), and near neurons had a mean *WBI* of -0.01 which was marginally less than zero (p = 0.0537). Attention had a significant effect on *WBI*, giving rise to a larger *WBI* when attention was directed to the near surface than the far surface across near neurons (p = 5.4×10^−9^), far neurons (p = 0.007), and all neurons combined (p=3×10^−10^) (Fig. 4A).

**Figure 4.**
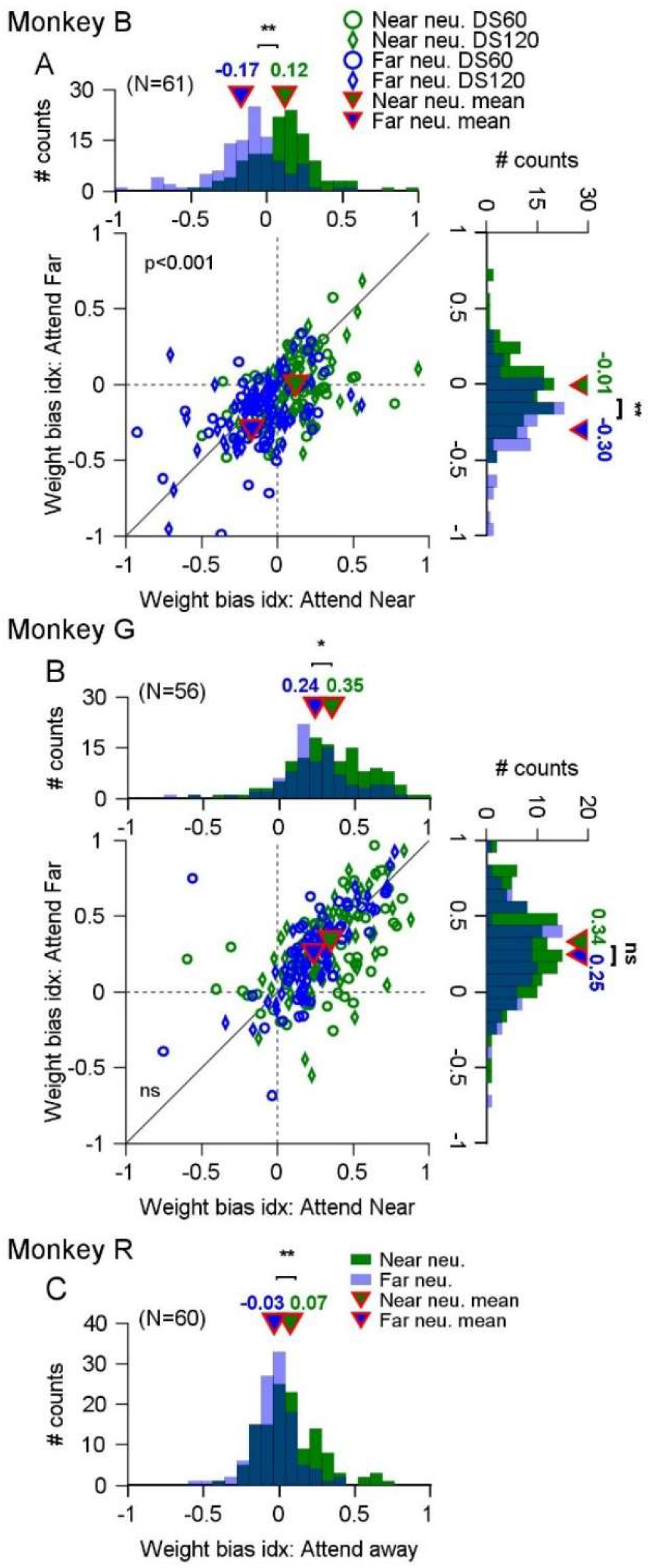
Disparity bias of near and far neurons and the effect of attention. **A. B**. Scatter plots and marginal distributions of weight bias indices (*WBI*) of near (green) and far (blue) neurons under attend-near and attend-far conditions, for monkeys B (**A**) and G (**B**). In the scatter plots, each point represents the *WBI* based on one bi-directional tuning curve from one neuron. Each neuron contributed to four points, from direction separations of 60° (circle) and 120° (diamond), and stimulus configurations of Near C/Far CC and Near CC/Far C. The *p*-value in the scatter plot was from a paired signed-rank test. The histograms show the marginal distributions of *WBI* in the attend-near (top) and attend-far (right) conditions, for near (green) and far (blue) neurons. **C**. The distributions of *WBI* for monkey R when attention was directed away from the RFs, for near (green) and far (blue) neurons. In all histograms, (**) indicates p < 0.001, (*) indicates p < 0.01, (ns) indicates p > 0.05 (one-sided signed-rank test). The triangles show the mean *WBI* values for near and far neurons. The p value (A) and ns (B) in the scatter plots indicate significance level of comparing *WBI* between attending near and attending far conditions for all neurons combined.

For monkey G, both near and far neurons showed a significant near bias, regardless of whether the near surface or the far surface was cued for attention (*WBI* significantly greater than 0, p = 6.09×10^−56^). Notably, near neurons showed a significantly larger *WBI* than the far neurons when attention was directed to the near surface (one-sided rank sum test, p < 0.01) and when the attending near and attending far trials were pooled (one-sided rank sum test, p = 0.002). For monkey G, attention did not change *WBI* significantly across the near neurons (one-sided signed-rank test, p = 0.29), the far neurons (p = 0.59), and all neurons combined (p = 0.41).

For monkey R, we found a similar result that the near neurons showed a significantly larger *WBI* than the far neurons, when attention was directed away from the RFs (one-sided rank sum test, p= 3.94×10^−5^) (Fig. 4C). Near neurons showed a significant near bias (mean *WBI* = 0.07, p = 4.3×10^−4^) and far neurons showed a significant far bias (mean *WBI* = -0.03, p = 0.003). In summary, MT neurons in monkeys B and R showed a significant bias toward the stimulus component moving at the more preferred disparity of the neurons. Although neurons in monkey G showed a predominant near bias, near neurons tended to show a stronger near bias than the far neurons.

### Relationship between disparity bias in response to two surfaces and disparity preference to a single surface

We asked whether a neuron’s disparity bias toward one depth of two overlapping surfaces was related to the neuron’s disparity preference to a single surface. To quantify the disparity bias toward one depth of two surfaces, we defined a two-surface disparity bias index (*TDBI*) (Eq. 3).

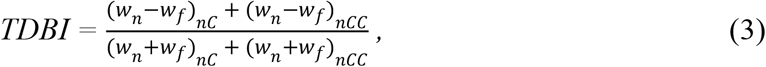

*TDBI* was similar to the weight bias index (*WBI*) (Eq. 2), except that *TBI* considered both stimulus configurations of Near C/Far CC and Near CC/Far C, and pooled the response weights obtained when the near component moved in the clockwise (*nC*) and counterclockwise (*nCC*) side of two motion directions. By doing so, *TDBI* balanced the weight difference between the near and far components contributed by potential directional bias, and therefore reflected the disparity bias. A positive *TDBI* indicates a disparity bias toward the near component of two surfaces and a negative *TDBI* indicates a bias toward the far component.

We also quantified a neuron’s preference for the disparity of a single surface, measured by a single-surface disparity preference index (*SDPI*):

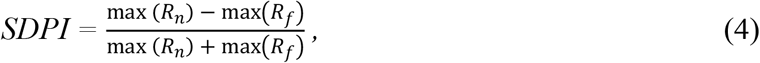

in which *R*_*n*_ and *R*_*f*_ were the direction tuning curves to the single near and far surfaces, respectively, when they were presented alone. The data samples from monkeys B and R were roughly balanced between near and far neurons. The neurons from monkeys B and R had median *SDPIs* of 0.0095 and 0.0045, respectively, which were not significantly different from zero (p > 0.24). In contrast, monkey G’s sample had more near neurons, with a median *SDPI* of 0.016, which was significantly positive (p = 0.025).

Figure 5 shows the relationship between the *TDBI* and *SDPI*. For all three animals, the *TDBI* was significantly correlated with the *SDPI* (Pearson correlation coefficient *r* = 0.57, 0.31, and 0.24 for monkeys B, G, and R, respectively, p < 0.01). The positive correlation was robust regardless of whether monkeys B and G were cued to attend the near surface or the far surface (Fig. 5A, B), or whether attention was directed away from the RFs (Fig. 5C). For monkey G, even though most of the neurons showed a bias toward the near disparity of the two surfaces, the positive correlation between the *TDBI* and *SDPI* still holds. These results suggest a common trend across the animals that the disparity bias of a neuron in response to two surfaces is positively correlated with the disparity preference of the neuron to a single surface.

**Figure 5.**
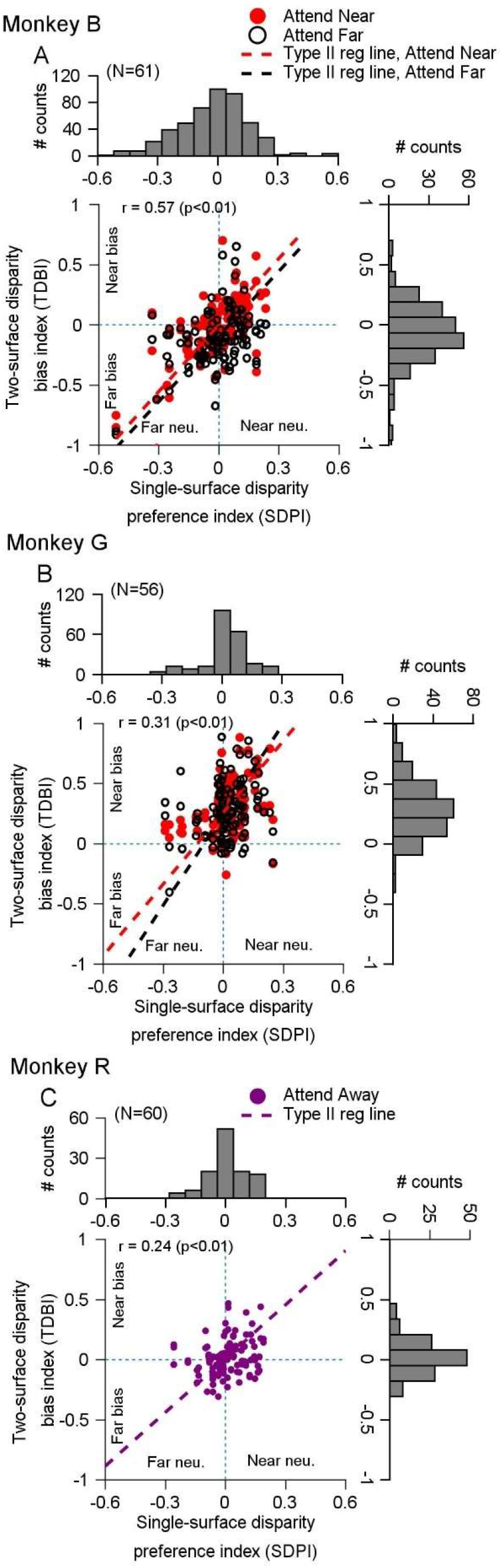
Relationship between the disparity bias to two surfaces and the disparity preference for a single surface. **A**. Scatter plot showing two-surface disparity bias index (*TDBI*) versus single-surface disparity preference index (*SDPI*) for each neuron from monkey B. Solid red circles represent attending near conditions, and open black circles represent attending far conditions. Each neuron contributes four data points (60° and 120° direction separations under both attention conditions). Dashed lines show Type II regression fits for attending near (red) and far (black) conditions. The Pearson correlation coefficient (*r*) is based on pooled results. Histograms display the distributions of *TDBI* (right) and *SDPI* (top). **B**. Results from monkey G. **C**. Results from monkey R, with attention directed away from the RFs.

### Timecourse of the neuronal response tuning to bi-directional stimuli moving at different depths

To better understand the nature of disparity bias and its underlying mechanisms, we analyzed the timecourse of response tuning curves to bi-directional stimuli at different depths. We aimed to address three key questions. First, does disparity bias emerge immediately at the start of the neural response or develop over time? Second, what is the relationship between disparity bias and attention modulation? Third, across neurons preferring near and far disparities, does the MT neuron population exhibit an overall bias towards the near or far disparity of the two surfaces?

Figure 6 shows the population-averaged tuning curves of monkey B to the bi-directional stimuli, calculated within a time window of 50 ms and sliding at a step of 10 ms. As in Figure 3, we combined the conditions in which the near component moved at either the clockwise or the counterclockwise side of the two directions and plotted the near component at the clockwise side of the two motion directions. As such, at positive VA directions, the near component moved in a more preferred direction of the neurons and elicited a stronger response than the far component (Fig. 6A1-D1). Specifically, at the VA direction of 30⁰ and 60⁰, the near component moved in the PD (0⁰) of the neuron when the direction separation was 60⁰ and 120⁰, respectively. We referred to VA 30⁰ or 60⁰ as “Near PD” for direction separation of 60⁰ and 120⁰, respectively. Conversely, at negative VA directions, the far component moved in a more preferred direction and elicited a stronger response (Figs. 6A1-D1). At the VA direction of -30⁰ and -60⁰, the far component moved in the PD (0⁰) at the direction separation of 60⁰ and 120⁰, respectively. We referred to VA -30⁰ or -60⁰ as “Far PD”. When the response tuning to the bi-directional stimuli is biased toward the near component, the tuning curve should skew to the positive VA directions. Likewise, when the bias is toward the far component, the tuning should skew to the negative VA directions.

**Figure 6.**
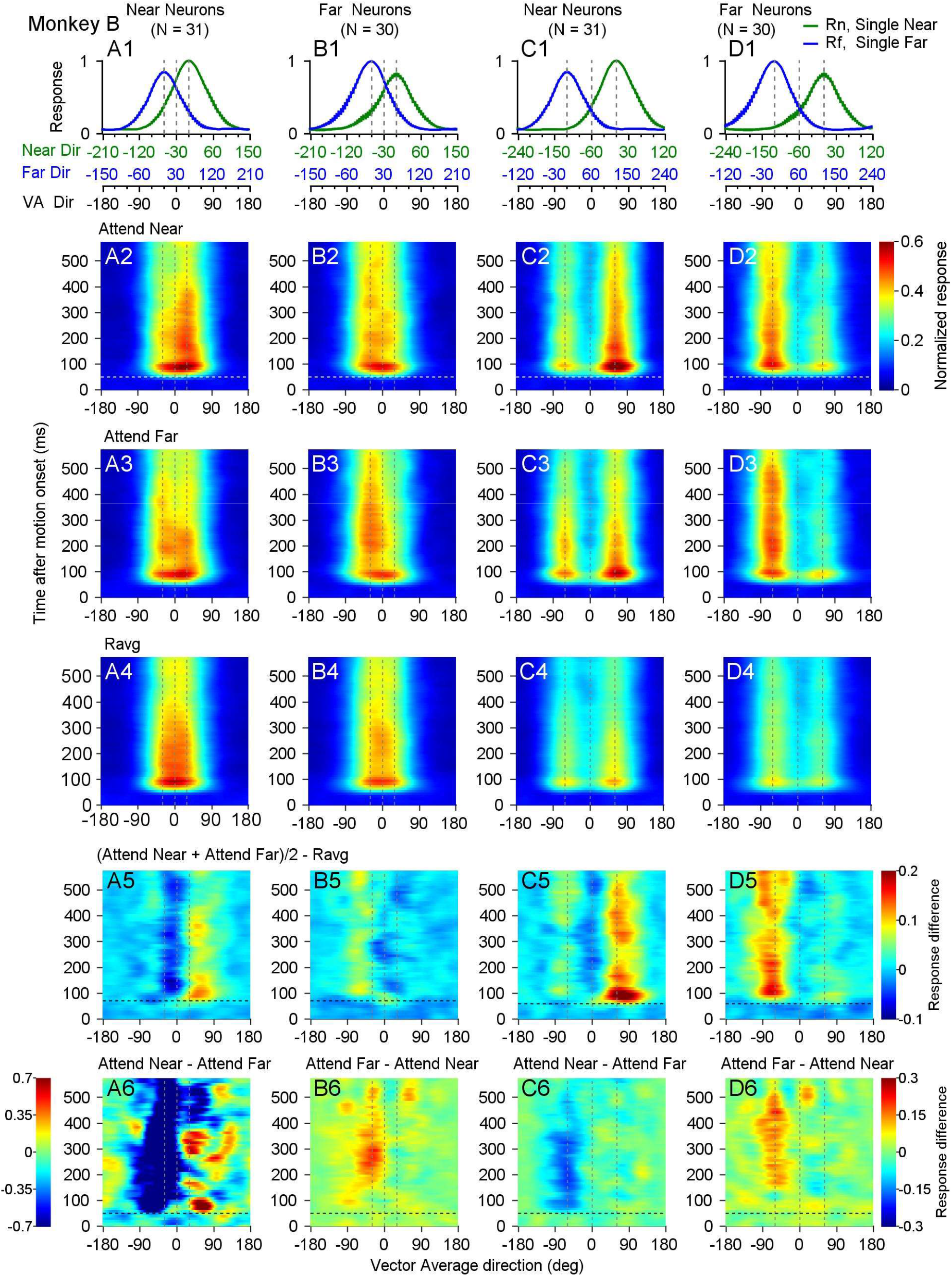
Timecourse of response tuning to bi-directional stimuli moving at different depths in monkey B. **Columns A, B**. Responses to 60° direction separation. **Columns C, D**. Responses to 120° direction separation. **Columns A, C**. Near neurons. **Columns B, D**. Far neurons. **A1-D1**. Population-averaged response tuning curves to single component directions at near (green) or far (blue) depth. The width of the curve for *R*_*n*_, *R*_*f*_ represents the standard error. Vertical dashed lines mark the VA direction at the neuron’s PD (0°) (middle line), and when far (left line) or near (right line) components move in the PD. **A2-D2**. Timecourse of population-averaged tuning curves to bi-directional stimuli under Attend Near condition. Horizontal dashed lines at 50 ms after motion onset roughly mark the response onset. **A3-D3**. Responses under Attend Far condition. **A4-D4**. Average responses to near and far components (*Ravg*). **A5-D5**. Average responses under Attend Near and Attend Far conditions, subtracting *Ravg*. Horizontal dashed lines at 70 ms (A5) roughly indicate the onset of near bias at 60° direction separation. Horizontal dashed lines at 60 ms (C5) roughly indicate the onset of near bias at 120° direction separation. The far bias of far neurons is later than the near bias of near neurons (B5, D5). **A6-D6**. Difference between responses under two attention conditions. A6, C6. Attend Near – Attend Far. B6, D6. Attend Far – Attend Near. Horizontal dashed lines at 50 ms (A6-D6) roughly indicate the onset of attention effect for near neurons (A6, C6). The effect of attention for far neurons is later than for near neurons (B6, D6).

At 60⁰ DS, the near neurons showed a near bias slightly delayed for about 10-20 ms relative to the neural response (Fig. 6A2, A3, Supplementary Fig. 1A). Attending the near surface made the near bias stronger and more persistent over time (Fig. 6A2), in comparison to attending the far surface (Fig. 6A3). Attending the far surface did not abolish the initial near bias, but later shifted the tuning back and forth between the near and far components and made the tuning broader than attending near (Fig. 6A3). The far neurons showed a far bias delayed about 40 ms relative to the neural response (Fig. 6B2, B3, Supplementary Figure SFig. 1C). The far bias of the far neurons appeared later than the near bias of the near neurons. Attending the far surface made the far bias more robust and persistent over time (Fig. 6B3) than attending near (Fig. 6B2). For both the near and far neurons, the average of the responses to the near and far components (*Ravg*) was roughly balanced around VA 0⁰, with a slight near (Fig. 6A4) and far (Fig. 6B4) skewness, respectively, due to the difference in the responses to the near and far components.

Similarly, at 120⁰ DS, the near neurons showed a near bias (Fig. 6C2, C3), and the far neurons showed a far bias (Fig. 6D2, D3). The disparity bias appeared sooner than that at 60⁰ DS and (SFig. 1B vs. 1A, 1D vs. 1C), again, the near bias of the near neurons emerged sooner than the far bias of the far neurons (SFig. 1B vs. 1D). Selective attention modulated the responses but was not sufficient to change the disparity bias according to attention (Fig. 6C2, C3, D2, D3). For both the near and far neurons, the *Ravg* was slightly skewed toward the neurons’ preferred disparity (Fig. 6C4, 6D4).

To separate the disparity bias from the effect of selective attention, we averaged the tuning curves obtained from “Attend Near” and “Attend Far” conditions. If the effects of attending to the near and far surfaces on the direction tuning curve of the same neuron have similar magnitude and timecourse, averaging the two would neutralize the impact of attention modulation. Consequently, any remaining bias in the tuning curve would be attributed to disparity bias. To illustrate how the recorded response tuning to the bi-directional stimuli differed from the average of the component responses over time, we subtracted *Ravg* from the attention-averaged responses (Fig. 6A5-D5).

At both 60⁰ and 120⁰ DS, we found that the near neurons showed a sooner, initially stronger and more transient near bias (Fig. 6A5, C5) than the far bias of the far neurons (Fig. 6B5, D5). Furthermore, the near bias at 60⁰ DS was slightly delayed relative to that at 120⁰ DS (comparing Fig. 6A5 with C5, and Fig. 6B5 with D5, SFig. 1A with 1B). Relative to response averaging, at 60⁰ DS, the disparity bias was reflected in response facilitation at the biased side, and suppression near VA 0⁰ and the other side of the VA directions (Fig. 6A5, B5). At 120⁰ DS, the disparity bias was reflected in stronger response facilitation on the biased side than on the other side (Fig. 6C5, D5). The suppression near VA 0⁰ was stronger for the near neurons than the far neurons (comparing Fig. 6A5 vs B5, and C5 vs D5). Interestingly, at 60⁰ DS, the far neurons initially showed a near bias, relative to the average of the component responses, for about 40 ms before changing to the far bias (Fig. 6B5 and the thin purple curve in SFig. 1C).

To examine the effects of selective attention, we subtracted the response tuning of the attending-far condition from the attending-near condition for the near neurons (Fig. 6A6, C6), and subtracted the attending-near condition from the attending-far condition for the far neurons (Fig. 6B6, D6). For near neurons, the attention modulation was reflected in response facilitation at positive VA directions around “Near PD” and/or suppression at negative VA directions around “Far PD” (Fig. 6A6, C6). The facilitation (Fig. 6A6) and suppression (Fig. 6A6, C6) occurred soon after the response onset. In contrast, for far neurons, the attention modulation was mainly reflected in much-delayed facilitation at negative VA directions around “Far PD”, without suppression at the positive VA directions around “Near PD” (Fig. 6B6, D6).

In summary, the timecourse analysis revealed several findings from monkey B. First, the disparity bias is often delayed relative to the response onset. The delay is slightly longer at the smaller (60⁰) than larger (120⁰) DS, likely reflecting a longer processing time needed for segmentation at smaller DS. Second, the near bias of the near neuron emerges sooner than the far bias of the far neuron. Third, the disparity bias is not the result of attention modulation but attention can modulate the disparity bias. Attention modulation for the near neurons also occurs sooner than the far neurons (comparing Fig. 6A6, C6 with B6, D6).

Figure 7 shows the timecourse of direction tuning from monkey G. At 60⁰ DS, the near neurons showed the development of near bias over a period of about 80 ms, starting from around 110 ms to fully biased to the near side around 190 ms (Fig. 7A2, A3, A5, SFig. 2A). At 120⁰ DS, the near neurons showed a stronger near bias than at 60⁰ DS (Fig. 7C2, C3, C5, SFig. 2B). The near bias was stronger and more persistent when attention was directed to the near surface (Fig. 7A2, C2) than directed to the far surface (Fig. 7A3, C3). Far neurons of monkey G also showed a near bias (Fig. 7B2, B3, B5, D2, D3, D5, SFig. 2C, 2D), displaying a timecourse similar to but slightly later than that of the near neurons (SFig. 2). The responses of the far neurons at the positive VA directions around the “near PD” were robust and sustained throughout the motion period, making the tuning curve broader than the near neurons (comparing Fig. 7B2 with A2, and B3 with A3).

**Figure 7.**
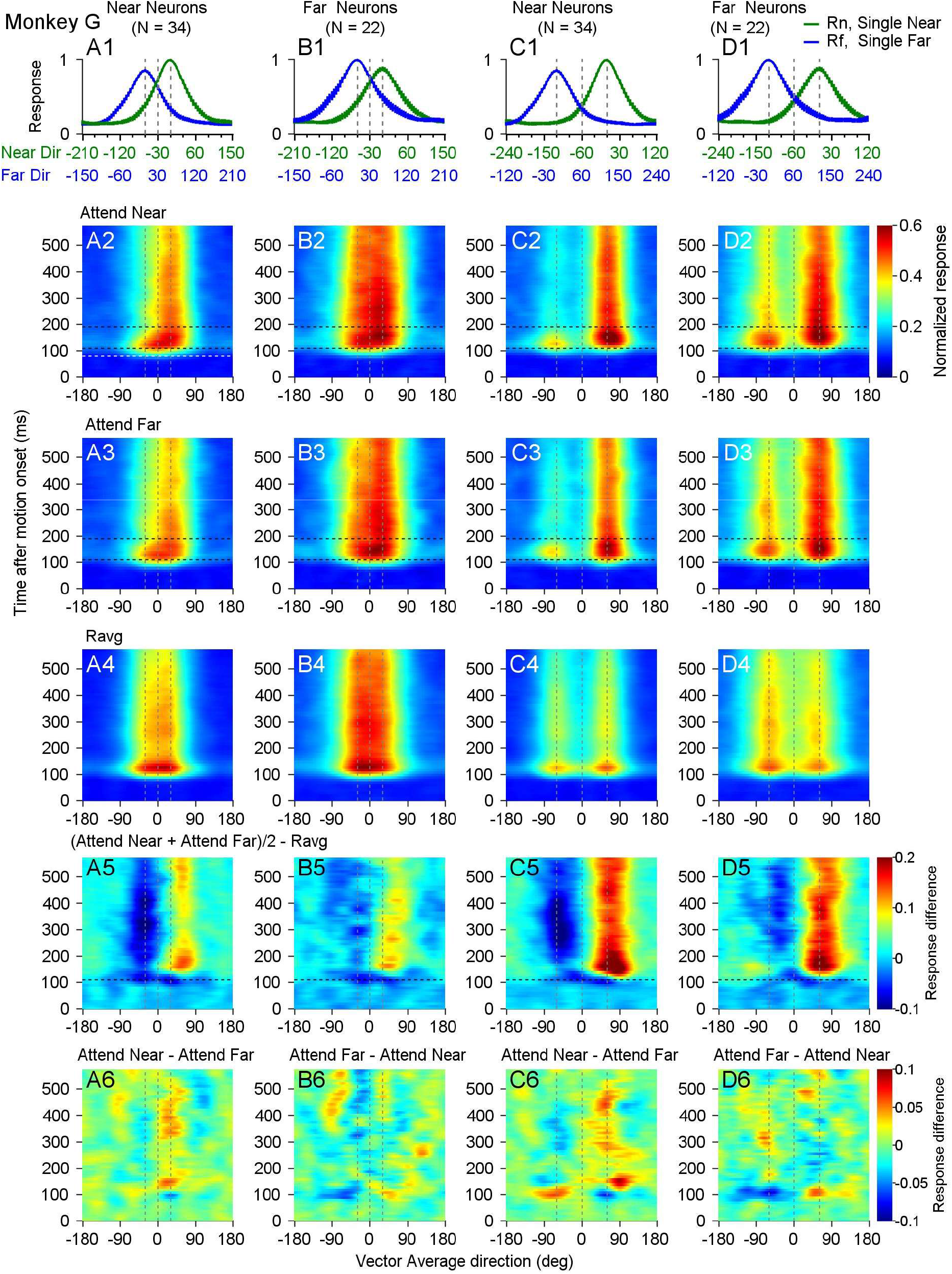
Timecourse of response tuning to bi-directional stimuli moving at different depths in monkey G. The convention follows that of Figure 6. **A1-D1**. Population-averaged response tuning curves to single component directions at near (green) or far (blue) depth. **A2-D2**. Timecourse of population-averaged tuning curves to bi-directional stimuli under Attend Near condition. In A2, the white horizontal dashed line at 80 ms indicates response onset and the two black dashed lines at 110 ms and 190 ms indicate the time period during which the near bias of near neurons develops. **A3-D3**. Responses under Attend Far condition. The black dashed lines at 110 ms and 190 ms are also plotted in B2-D2 and A3-D3 for reference. **A4-D4**. Average responses to near and far components (*Ravg*). **A5-D5**. Average responses under Attend Near and Attend Far conditions, subtracting *Ravg*. The black dashed lines are at 110 ms as a reference. **A6-D6**. Difference between responses under two attention conditions. Vertical dashed lines in all panels mark the VA direction at the neuron’s PD (0°) (middle line), the far PD (left line), and the near PD (right line).

Although the weight analysis based on the full tuning curve and the entire motion response period did not reveal a significant attention effect for monkey G (Fig. 4B), the timecourse analysis indicated a small effect of attention modulation, which was more evident in the near neurons than in the far neurons. The attention modulation for the near neurons mainly reflected in response facilitation at the positive VA directions around the “near PD” when the response of attending far was subtracted from attending near (Fig. 7A6, C6). For near neurons, there was small and brief facilitation toward negative VA directions around the far PD at the beginning of neural response (about 100 ms after motion onset) (Fig. 7A6, C6), suggesting that, in the attending-near condition, monkey G may have initially attended to the far surface briefly before attending to the cued near surface.

Far neurons showed little attention modulation at 60⁰ DS (Fig. 7B6), but showed small and delayed facilitation at the negative VA directions around the “far PD” at 120⁰ DS when the response tuning of attending near was subtracted from attending far (Fig. 7D6). Similar to the near neurons, the far neurons showed a small initial facilitation toward positive VA directions around the near PD at about 100 ms (Fig. 7B6, D6), suggesting that, in the attending-far condition, monkey G may have attended to the near surface briefly before attending to the cued far surface. Nevertheless, regardless of the effect of attention modulation, the near bias was robust throughout the motion response period in both the near and far neurons (Fig. 7A2-D2, A3-D3, A5-D5).

Figure 8 shows the timecourse of the response tuning to bi-directional stimuli at different depths in monkey R, with the animal’s attention directed away from the RFs. The near and far neurons showed disparity bias toward their preferred disparity (Fig. 8A2-D2, A4-D4). The near bias of the near neurons appeared sooner than the far bias of the far neurons (Fig. 8A4-D4, SFig. 3). For the far neurons, the early responses were roughly balanced between VA directions around the far PD and near PD, and the far bias only became evident 160 ms and 100 ms after motion onset for DS60⁰ and 120⁰, respectively (Fig. 8B4, D4, SFig. 3C, D). As found in monkey B, the far neurons from monkey R initially showed a near bias relative to the average of the component responses, before gradually changing to the far bias (thin purple curves in SFig. 3C, 3D).

**Figure 8.**
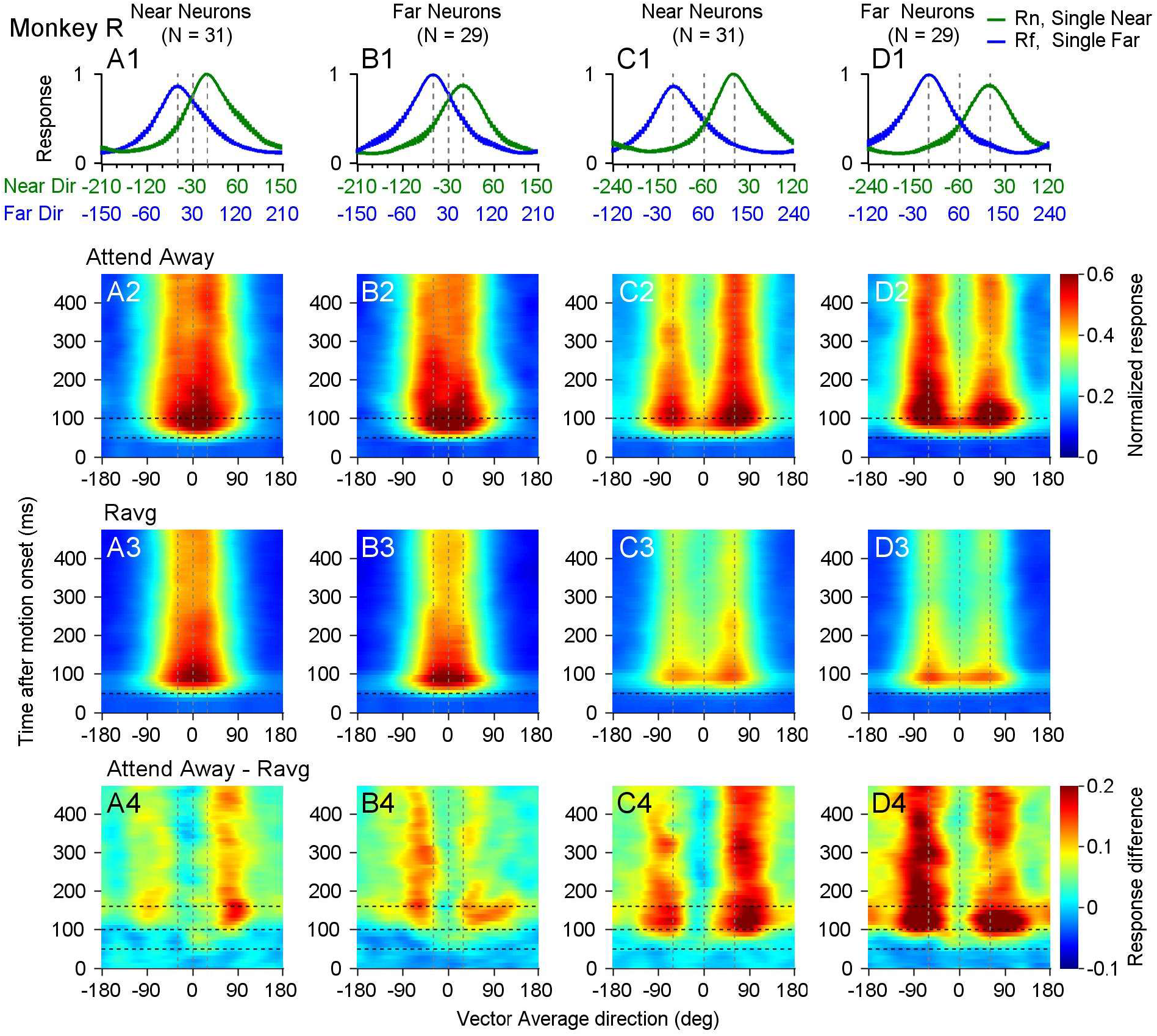
Timecourse of response tuning to bi-directional stimuli moving at different depths in monkey R. The convention is similar to that of Figures 6 and 7. **A1-D1**. Population-averaged response tuning curves to single component directions at near (green) or far (blue) depth. **A2-D2**. Timecourse of population-averaged tuning curves to bi-directional stimuli with the animal’s attention directed away from the RFs. The two horizontal dashed lines are at 50 ms and 100 ms after motion onset as refenerences. **A3-D3**. Average responses to near and far components (*Ravg*). The horizontal dashed line is at 50 ms. **A4-D4**. Responses to bi-directional stimuli subtracting *Ravg*. The three horizontal dashed lines are at 50 ms, 100 ms, and 160 ms after motion onset. Vertical dashed lines in all panels mark the VA PD (middle line), the far PD (left line), and the near PD (right line).

So far, we have characterized the timecourse of disparity bias separately for near and far neurons. Next, we examined whether MT neurons as a whole showed an overall bias toward the near or far disparity of overlapping surfaces over time. We averaged the tuning curves to the bidirectional stimuli within a sliding time window of 50 ms across near and far neurons for each animal. To assess disparity bias, we subtracted *Ravg* from the average Attend-Near and Attend-Far responses for monkeys B (Fig. 9A1, B1) and G (Fig. 9A2, B2), and from the attention-away responses for monkey R (Fig. 9A3, B3). During the early response, all three animals exhibited a near bias for both 60⁰ DS (Fig. 9A1-A3) and 120⁰ DS (Fig. 9B1-B3), with stronger responses at positive VA directions than at negative VA directions.

**Figure 9.**
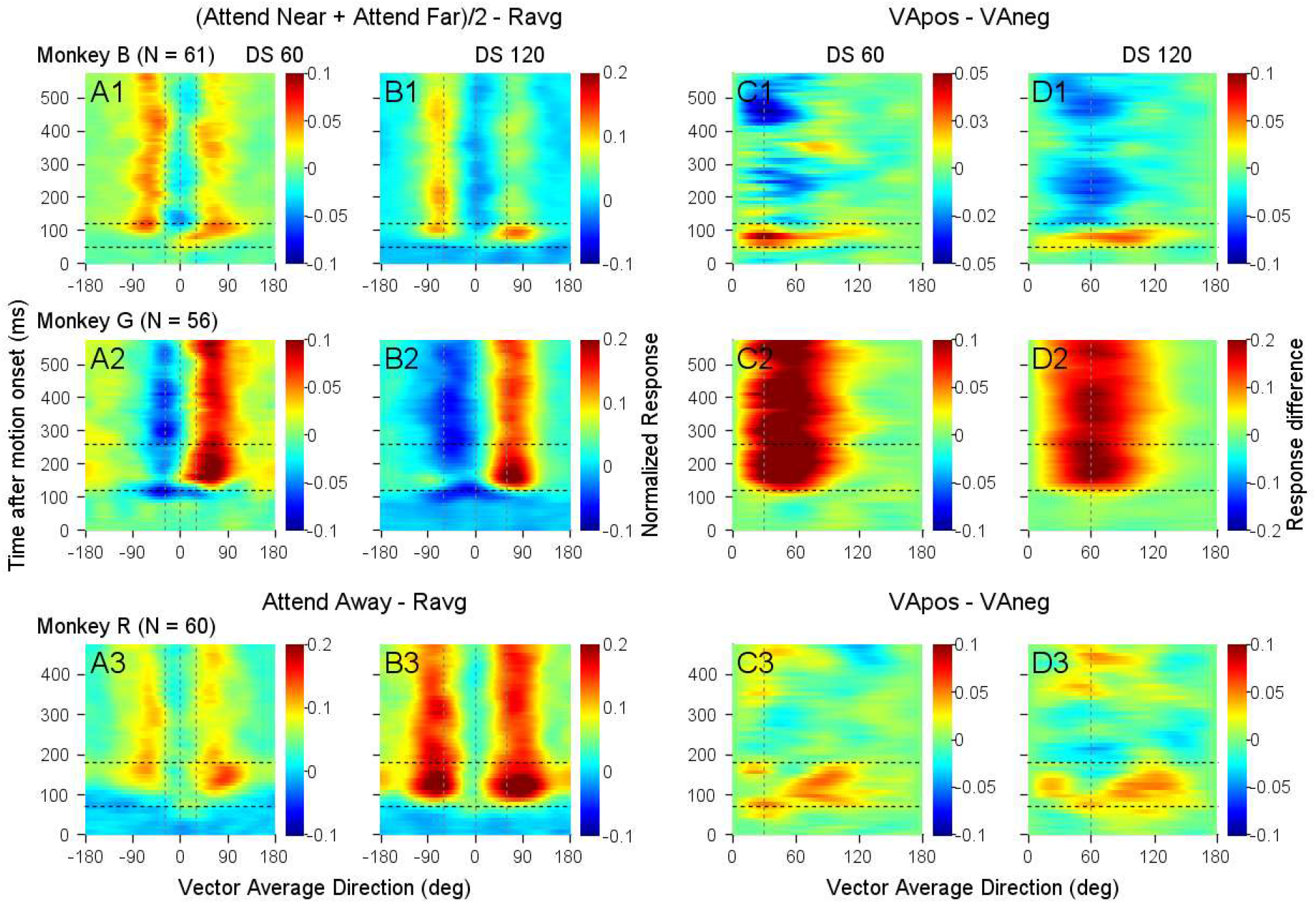
Early near bias across all neurons in three monkeys. **A1-D1**. Responses from monkey B. Horizontal dashed lines indicate 50 and 120 ms after motion onset. **A2-D2**. Responses from monkey G. Horizontal lines indicate 120 and 260 ms. **A3-D3**. Responses from monkey R. Horizontal lines indicate 70 and 180 ms. **Columns A, C**. Responses to 60° DS. **Columns B, D**. Responses to 120° DS. **A1, B1, A2, B2**. Average responses under Attend Near and Attend Far conditions, subtracting Ravg. **A3, B3**. Responses to bi-directional stimuli with attention directed away from the RFs, subtracting Ravg. Vertical dashed lines in columns A and B mark VA PD (middle line), far PD (left line), and near PD (right line). **Columns C, D**. Responses at negative VA directions were flipped relative to VA 0° and subtracted from responses at positive VA directions. The vertical dashed line indicates the component PD.

To quantify the disparity bias, we flipped the responses at negative VA directions relative to VA 0° and subtracted them from the responses at positive VA directions (Fig. 9C1-C3 and D1-D3). Positive values of the response difference (shown in black-red-yellow) indicate a near bias, while negative values (shown in blue-cyan) indicate a far bias. For all three animals, the response differences in the early time windows (integrated within VA directions ranging from 15⁰ to 120⁰ in Fig. 9C1-C3, D1-D3) were significantly positive (p < 0.01 for monkeys B and G, and p ≤ 0.05 for monkey R, signed-rank test), indicating a near bias (Table 1).

**Table 1.** Time period showing significant near bias.

| **Time after motion onset** | Mid-point of 50-ms time bins (Time period measured in 10-ms bin)<br>(p value) |  |
| --- | --- | --- |
| **Direction separation** | **DS 60°** | **DS 120°** |
| Monkey B (N = 61) | 55 – 85 ms (30 – 110 ms)<br>($p < 0.01$) | 45 – 85 ms (20 – 110 ms)<br>($p < 0.01$) |
| Monkey G (N = 56) | 115 – 575 ms (90 – 600 ms)<br>($p < 0.01$) | 105 – 575 ms (80 – 600 ms)<br>($p < 0.01$) |
| Monkey R (N = 60) | 35 – 55 ms (10 – 80 ms)<br>($p < 0.05$) | 75 ms (50 – 100 ms)<br>($p = 0.05$) |

Whether the population-averaged response tuning showed a near or a far bias depended on the proportion of near and far neurons in the data sample and their disparity preferences as measured by *SDPI*. As shown in Figure 5, data samples from monkeys B and R were roughly balanced between near and far neurons. Therefore, the early near bias observed in the population-averaged tuning curves of monkeys B and R (Fig. 9C1, D1, C3, D3) cannot be attributed to the incidental disparity preferences of the neurons in the data samples. Our findings indicate that near neurons showed an earlier near bias than far neurons showing far bias in monkeys B (Fig. 6, SFig. 1) and R (Fig. 8, SFig. 3), likely contributing to the early near bias observed in the overall population. While monkey G’s sample had more near neurons than far neurons, the consistent near bias found in both early and late responses (Fig. 9C2, D2) is due to both near and far neurons exhibiting the near bias (Fig. 7, SFig. 2).

### Attention modulation of neuronal response is object-based

Figure 4 describes the effect of attention on the full response tuning curve to bi-directional stimuli, as measured by the impact on the weights for the near and far components. We further asked whether the effect of attention was feature-based or object-based when the animals were cued to attend a stereoscopic depth of a surface. If the attention was feature-based (disparity-based), we expected to find that when attention was directed to a more preferred disparity of a neuron, the neuronal response would be enhanced regardless of the motion direction of the surface. On the other hand, if attention was object-based (surface-based), we expected to find that when attention was directed to a surface that elicited a stronger response than the other surface, the neuronal response would be enhanced regardless of whether the attended surface was located at the more or less preferred disparity of the neuron.

To answer this question, we measured the effect of attention directed to a stimulus component that was located at either the more preferred or less preferred disparity and moving in either the more preferred or less preferred direction of the neuron. We quantified the effect using two attention modulation indices (AMI). One index *AMI_disparity* would be positive across conditions if the effect of attention was feature-based (Eq. 5). Whereas the other index *AMI_surface* would be positive if the effect of attention was object-based (Eq. 6).

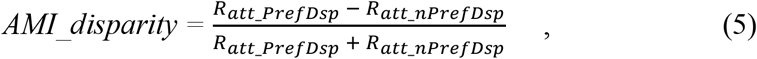

in which *R*_*att_PrefDsp*_ and *R*_*att_nPrefDsp*_ were the responses to the overlapping stimuli when attention was directed to the more preferred disparity and less preferred disparity of the neuron, respectively.

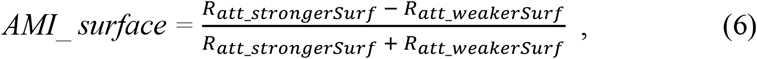

in which *R*_*att_strongerSurf*_ and *R*_*att_weakerSurf*_ were the responses to the overlapping stimuli when attention was directed to a surface that elicited a stronger response and a weaker response of the neuron, respectively.

We found that the results from monkey B were consistent with object-based attention. Figure 10A and B showed the response tuning curves of two representative neurons from monkey B, one neuron preferred the near disparity (Fig. 10A) and the other preferred the far disparity (Fig. 10B). When attention was directed to a surface that elicited a stronger component response, regardless of whether the surface was located at the more preferred or less preferred disparity of the neuron, the response was always enhanced relative to attending the other surface (Fig. 10A, 10B). This trend was consistent across the neuronal population as captured by always positive *AMI_surface* values (Fig. 10C) for the near and far neurons, and for 60° and 120° DSs, respectively.

**Figure 10.**
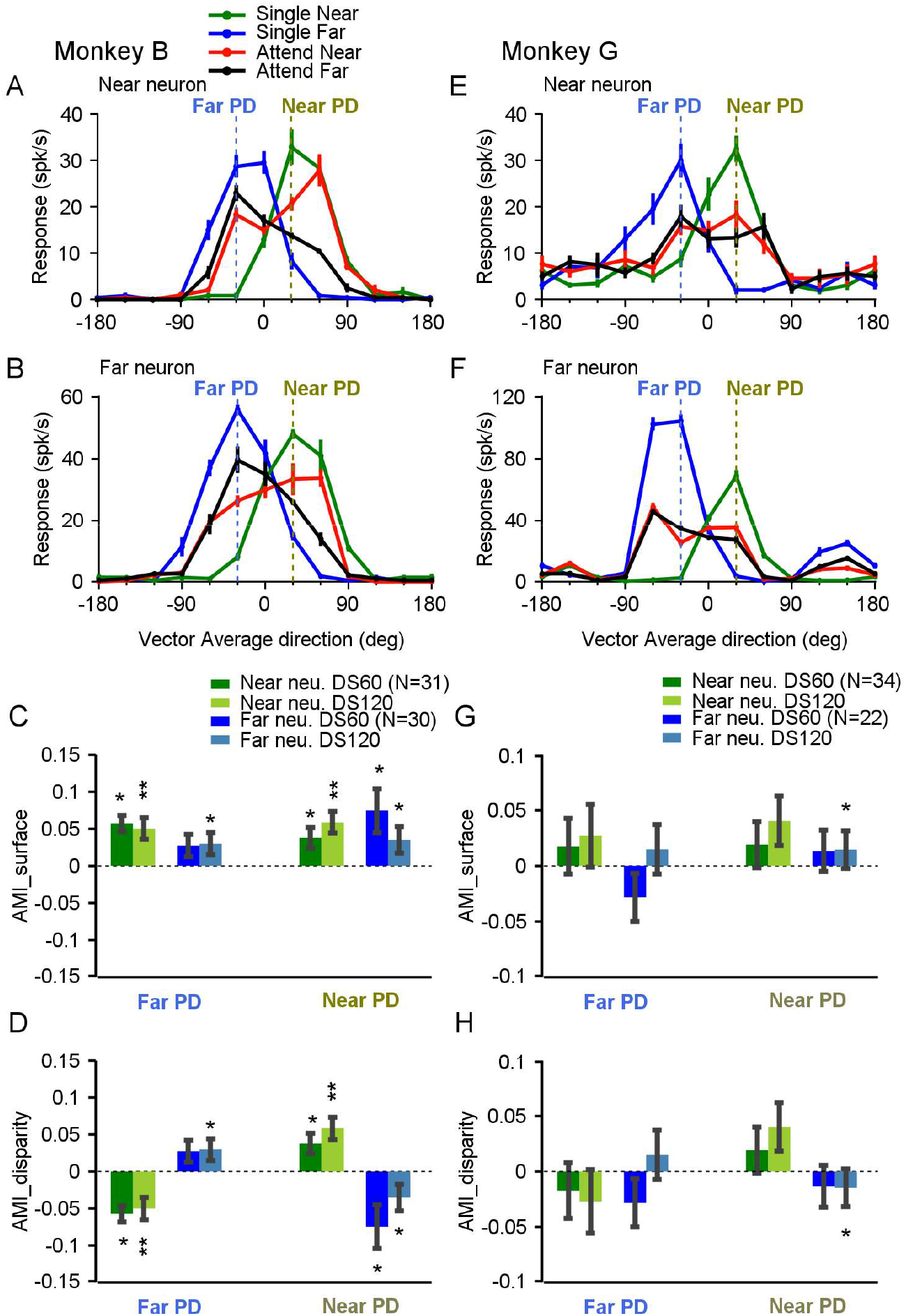
Attention modulation of neuronal responses to bi-directional stimuli moving at two depths for monkey B (A-D) and monkey G (E-H). **A, B, E, F**. Tuning curves of four example neurons to bi-directional stimuli. The convention follows that of Figure 2. Error bars represent the standard error. The vertical dashed lines indicate the Near PD and the Far PD. **C, G**. Population-averaged object-based attention modulation indices (*AMI_surface*) were calculated separately using responses to bi-directional stimuli at the Far PD and the Near PD, across near neurons and far neurons, and for 60° and 120° DSs. **D, H**. Disparity-based attention modulation indices (*AMI_disparity*). Responses of monkey B were calculated from 0 to 600 ms following the motion onset. Responses of monkey G were calculated from 110 to 410 ms. (**) indicates p < 0.01, (*) indicates p < 0.05, and no asterisk indicates p > 0.05 (one-sided signed-rank test to compare *AMI* with zero).

In contrast, when attention was directed to the surface located at the more preferred disparity of the neuron (near in Fig. 10A and far in 10B) and the surface moved in a less preferred direction, resulting in a weaker component response, the response to both surfaces was reduced relative to attending to the other surface (the red curve at “Far PD” in Fig. 10A and the black curve at “Near PD” in Fig. 10B). Furthermore, when attention was directed to the surface located at the less preferred disparity of the neuron (far in Fig. 10A and near in 10B) and the surface moved in a more preferred direction, resulting in a stronger component response, the response to both surfaces was nevertheless enhanced (the black curve at “far PD” in Fig. 10A and the red curve at “near PD” in Fig. 10B). In other words, the enhancement or reduction of the response by attention was not dependent on whether the attended surface was the more preferred or less preferred disparity of the neuron, respectively, but rather dependent on whether the attended surface elicted a stronger or weaker component response. As a result, *AMI_disparity* averaged across the neuronal population changed sign with stimulus conditions (Fig. 10D), suggesting that the effect of attention was not feature-based.

The attention effect found in monkey G was minimal when measured by the response weights based on the response during the whole stimulus motion period (Fig. 4B). However, the timecourse of the neuronal responses revealed that the attention effect in monkey G was visible in most conditions between 110 ms and 410 ms after motion onset (Fig. 7A6, C6, D6), except for far neurons responding to a 60⁰ DS (Fig. 7B6). Therefore, we used this time window to calculate direction tuning curves (Fig. 10E, F) and the *AMIs*. Although the attention effect in monkey G was not statistically significant for most conditions, except for the responses of the far neurons to 60⁰ DS, the trend of *AMI* in monkey G (Fig. 10G, H) was consistent with monkey B (Fig. 10C, D) and supported the idea of object-based attention modulation.

### Modeling neural responses to two surfaces at different depths using a unified model

Our neural data revealed that MT responses to overlapping moving stimuli separated in depth can be explained by a weighted sum of responses to individual stimulus components presented alone. These weights were influenced by the neuron’s disparity preference to single surfaces (Fig. 5) and the effects of selective attention (Fig. 4 and Fig. 10). While the general trend was consistent across animals, we also observed individual differences. Specifically, in monkey G, both near and far neurons exhibited a dominant near bias, though near neurons showed a stronger bias than far neurons (Fig. 5). In contrast, in monkey B, the near neurons showed a near bias and the far neurons showed a far bias. These findings prompted us to seek a unified model capable of explaining the seemingly complex relationships.

The divisive normalization model (Carandini and Heeger, 2012) has been used to explain a wide array of phenomena, including neuronal responses to multiple visual stimuli (Britten and Heuer, 1999; Heuer and Britten, 2002; Busse et al., 2009; Xiao et al., 2014; Xiao and Huang, 2015; Bao and Tsao, 2018; Wiesner et al., 2020; Huang et al., 2023) and the effects of attention (Reynolds and Heeger, 2009; Lee and Maunsell, 2009; Ni et al., 2012; Ni and Maunsell, 2019). While in the model, the division by the activity of a population of neurons (i.e., normalization) is well established, the factors contributing to the numerator are less clear. Typically, in response to multiple stimuli, the signal strength (such as luminance contrast or motion coherence) of each stimulus component is used in the numerator (e.g., Busse et al., 2009; Xiao et al., 2014). However, defining the signal strength of a sensory stimulus is not always straightforward (Wiesner et al., 2020). We have proposed that the weighting to a stimulus component of multiple stimuli is proportional to the activity of a population of neurons elicited by the same stimulus component (Xiao et al., 2014; Wiesner et al., 2020; Huang et al., 2023). We refer to this neuron population as the “weighting pool”. The nature and the scope of the “weighting pool” remain unclear.

Here, we expand on this idea and suggest that the size of the weighting pool can be variable. At one extreme, the weighting pool may encompass neurons with RFs at the same location but with all possible disparity preferences, a scenario we call “global pooling”. Conversely, at the other extreme, the weighting pool may consist solely of neurons with the same preferred disparity as the neuron under study, which we term “local pooling”. The size of the weighting pool can vary between these two extremes. Equations 7-10 describe the model without considering the effect of attention.

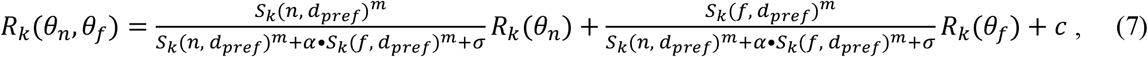

in which *R*_*k*_ *θ*_*n*_, *θ*_*f*_ is the model-predicted response of neuron *k* to two motion directions *θ*_*n*_ and *θ*_*f*_, moving at the near (*n)* and the far (*f*) disparity. *R*_*k*_*(θ*_*n*_) and *R*_*k*_*(θ*_*f*_) are the measured responses of the same neuron to the individual components *θ*_*n*_ and *θ*_*f*_, respectively. *m, α, σ*, and *c* are model parameters. *S*_*k*_ is the summed response of the weighting pool associated with neuron *k*, which has a preferred disparity *d*_*pref*_, to the disparity of a stimulus component. In our model, *S*_*k*_ is calculated by taking the dot product of a Gaussian window ***G***, centered on *d*_*pref*_, and the response of a population of neurons with all possible disparity preferences to the disparity of a stimulus component (Eq. 8).

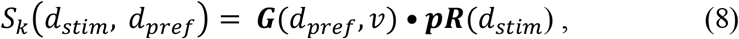

in which *d*_*stim*_ is the disparity of a stimulus component, ***pR*** is the response of a population of neurons with all possible disparity preferences (Eq. 9), and ***G*** is a Gaussian function with a standard deviation of *v* and centered on *d*_*pref*_ (Eq. 10).

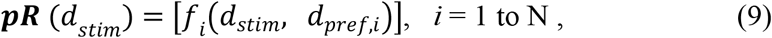

in which *f*_*i*_ is the disparity tuning function of neuron *i* in response to single disparities. *d*_*pref,i*_ is the preferred disparity of neuron *i*.

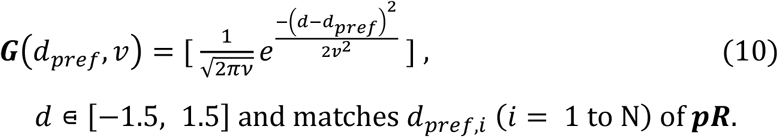

In other words, the extent of the weighting pool is controlled by the width of the Gaussian window ***G***. The parameter *v* is the standard deviation of ***G***. Both ***pR*** and ***G*** are vectors of the same length N. Given the relatively small number of neurons in our data set, the proportion of near and far neurons may not accurately reflect the true distribution of disparity preferences among MT neurons. Therefore, we used the disparity tuning data from a large sample of MT neurons (N = 479) as reported by DeAngelis and Uka (2003) to estimate ***pR***.

To model the effect of attention, we added the term *β* to equation 7. The model-predicted responses *R*_*k*_*(θ*_*na*_, *θ*_*f*_) and *R*_*k*_*(θ*_*n*_, *θ*_*fa*_) correspond to attention being directed to the near-surface and the far-surface, respectively (Eqs. 11, 12). The values of *β*_*An*_ and *β*_*Af*_ were derived from the weights of a linear weighted summation (LWS) model fit (see Methods and Eq. 16).

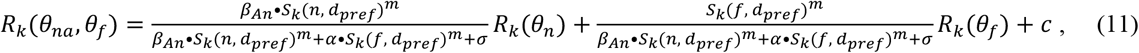

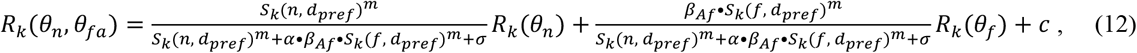

The ranges of the model parameters were constrained as follows: 0 < m < 100, c > 0, *σ* > 0, and *α* > 0. The parameter *α* scales the relative contributions of *S*_*k*_*(n)* and *S*_*k*_*(f)* to the denominator as in tuned normalization (Ni et al., 2012; Rust et al., 2006; Carandini et al., 1997).

Figure 11A illustrates the population neural response ***pR*** to a single near disparity of -0.1⁰ and a far disparity of 0.1⁰ of our visual stimuli, with the single stimulus moving in the preferred directions of the neurons. Figure 11A also displays examples of Gaussian function ***G*** with three different widths, centered on disparities of -0.5⁰ and 0.5⁰, demonstrating the transition from local pooling to global pooling. Due to the limited range of neurons’ preferred disparities in the data set, the Gaussian window is unavoidably truncated by the disparity borders. In practice, we found the area under the truncated Gaussian window, as a measure of the weighting pool size, that best fits the neuronal response using Equations 11 and 12.

**Figure 11.**
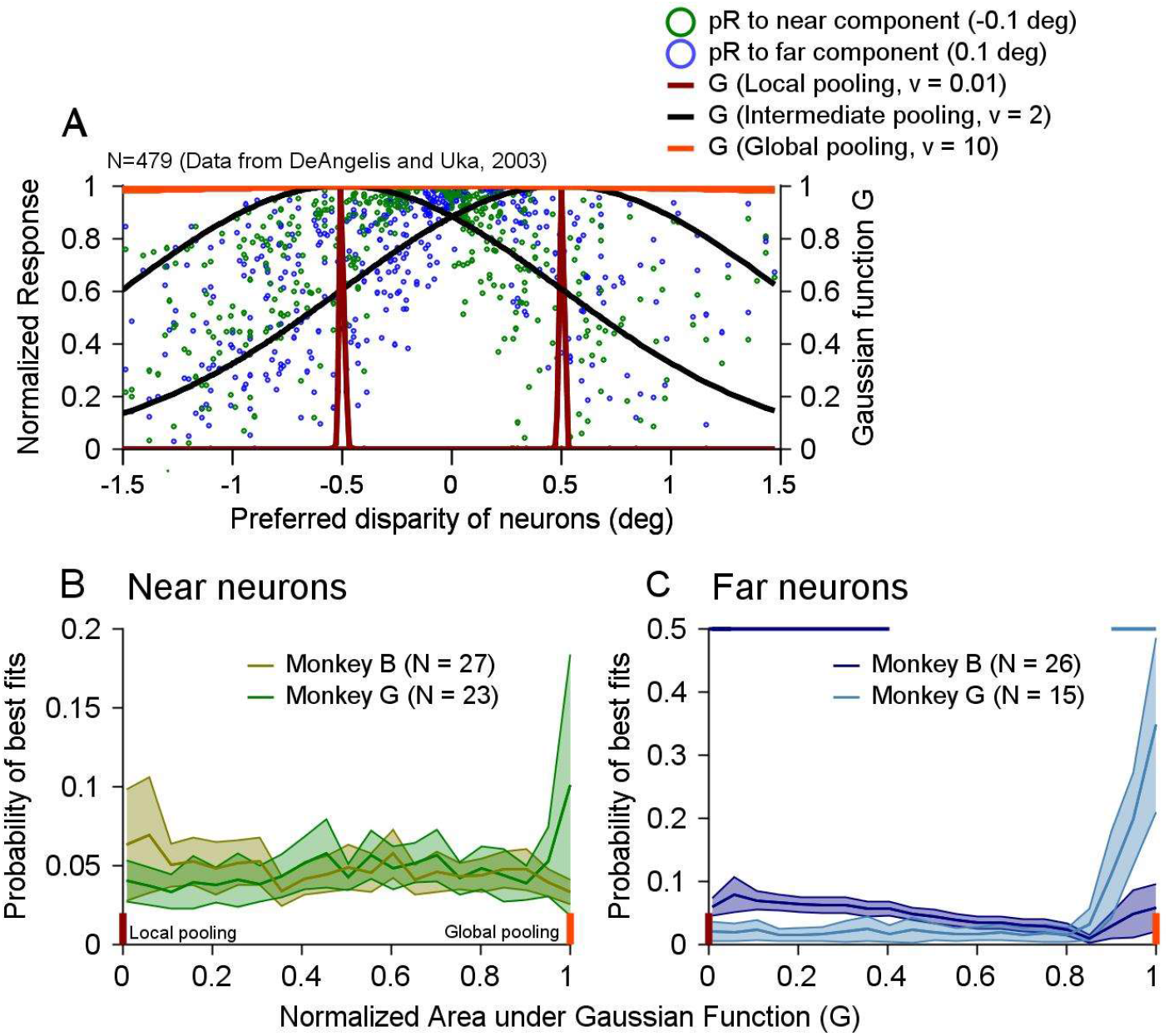
Modeling neural responses to bi-directional stimuli at two depths. **A**. MT population neural responses to component disparities of -0.1° and 0.1°, illustrating local, intermediate, and global pooling. The population responses are based on the disparity tuning curves of 479 MT neurons from DeAngelis and Uka (2003). *v* is the standard deviation of the Gaussian function, centered on the disparities of -0.5° and 0.5°, as two example *d*_*pref*_ of neurons under study. **B**. Distribution of pooling size providing the best fit for near neurons’ responses from monkeys B and G. **C**. Distribution for far neurons, with significant differences between monkeys B and G indicated by horizontal blue and cyan lines at those pooling sizes. Error bands show 95% confidence intervals measured by bootstrapping. Short vertical brown and red bars in B and C represent local and global pooling sizes. Only neurons for which complete disparity tuning curves were obtained were included in this analysis.

The model fitted the neuronal responses of monkeys B and G to the bi-directional stimuli reasonably well. For monkey B, the mean *PV* was 89 % (SD = 11.2), and for monkey G, it was 80% (SD = 17.5). We compared the weighting pool size that provided the best fit for the near neurons (Fig. 11B) and the far neurons (Fig. 11C) between monkeys B and G. For the near neurons, monkey B had more neurons that were best fitted with local pooling, while monkey G had more neurons that were best fitted with global pooling. However, these differences were not statistically significant as measured by 95% confidence intervals using bootstrapping (Fig. 11B).

For the far neurons, local pooling provided the best fit for significantly more neurons in monkey B than monkey G (at a normalized area of Gaussian window less than 0.4, p < 0.05), whereas global pooling provided the best fit for significantly more neurons in monkey G than monkey B (at area greater than 0.9, p < 0.05) (Fig. 11C). For 11 out of 15 far neurons (70%) from monkey G, the optimal area of the Gaussian window was greater than 0.85, indicating that global pooling best accounted for the responses of far neurons from monkey G. In contrast, local and intermediate pooling best accounted for the results from monkey B.

An intuitive explanation for these results is as follows. Because the overall MT population has more near neurons than far neurons (DeAngelis and Uka, 2003), the global pooling of MT population responses to -0.1° near disparity results in a slightly stronger response than to 0.1° far disparity. Consequently, the weight for the near component is greater than for the far component, irrespective of the disparity preference of the neuron under study. Conversely, the local pooling of MT population responses to the component disparities aligns with the disparity preference of the neuron under study. Thus, near neurons show a near bias, and far neurons show a far bias. Taken together, our findings demonstrate that a unified model with a variable size of the weighting pool and divisive normalization can explain both the disparity bias and the individual differences between the animals.

## Discussion

### Disparity bias and implication on the rules of representing multiple visual stimuli

Our laboratory previously found that when two overlapping random-dot stimuli move in different directions at the same depth, MT neurons exhibit a robust response bias toward the stimulus with higher motion coherence or luminance contrast (Xiao et al., 2014). Nearly all MT neurons respond more strongly to higher motion coherence (Britten et al., 1992, 1993; Xiao et al., 2014) and contrast (Cheng et al., 1994; Heuer and Britten, 2002; Sclar et al., 1990), showing consistent stimulus preference at both individual and population levels. However, different neurons prefer different values for features like binocular disparity. Our current study establishes the encoding rule for representing multiple stimuli when neurons have varying preferences for a stimulus feature.

We discovered that MT neurons displayed a disparity bias toward one of two overlapping surfaces at different depths. This bias positively correlated with each neuron’s preference for the disparity of a single surface when presented alone (Fig. 5). Despite individual differences among the animals, this positive correlation was consistent. Our finding may likely extend beyond binocular disparity. For instance, in another study from our lab, neuronal responses to two spatially separated stimuli moving in different directions showed a robust spatial bias, correlating with the spatial preferences of the neurons (Wiesner and Huang, 2019, SFN abstract). Thus, the disparity bias in this study may also be considered a “spatial bias,” aiding in segmenting multiple visual stimuli in 3D space.

### An extended normalization model and the neural mechanism underlying the disparity bias

In the standard divisive normalization model, the response to multiple stimuli is described by a weighted sum of the responses elicited by the individual stimulus components. The weight is proportional to the signal strength (such as contrast) of each stimulus component, normalized by a combination of the signal strengths of multiple stimuli (Busse et al., 2009; Carandini and Heeger, 2012; Ni et al., 2012; Xiao et al., 2014; Bao and Tsao, 2018). In terms of the neural implementation of the divisive normalization model, it has been proposed that the summed response of a population of neurons, referred to as the normalization pool, serves as the denominator in the normalization equation (Heeger, 1992; Carandini et al., 1997; Simoncelli and Heeger, 1998; Rust et al., 2006; Kouh and Poggio, 2008; Bao and Tsao, 2018; Wiesner et al., 2020). However, the neural implementation of the numerator in the normalization equation has been unclear when multiple visual stimuli are involved. We and other investigators have suggested that the summed response of a population of neurons elicited by a stimulus component may serve as the numerator and determine (together with the normalization) the weight of that component (Xiao and Huang, 2014; Bao and Tsao, 2018; Wiesner et al., 2020).

In this study, we propose a novel extension to the standard normalization model: the pool size of the neuron population in the numerator (the weighting pool) can vary. This variability allows the weighting for a stimulus component to depend on neurons with similar feature selectivity (local pooling) or those covering a full range of values (global pooling), or an intermediate range. With local pooling, a neuron’s response to two stimuli would bias toward its preferred feature value, as seen in monkeys B and R. Global pooling, however, reflects the preference of a full neuron population, leading to a near bias in response to two stimuli, regardless of individual neuron preferences. In the case of binocular disparity, MT neurons exhibit a higher proportion of neurons preferring near disparity over far disparity (DeAngelis and Uka, 2003). Therefore, global pooling results in a stronger response to near disparity. When the pool size is broad enough to represent this overall near preference but not entirely invariant to individual neuron preferences, a near bias and positive correlation with the neuron’s single-stimulus disparity preference emerges, as observed in monkey G.

Our model offers a unified explanation of the disparity bias observed across different animals. In the future, it would be interesting to understand what factors (such as experience, cognition, and other individual differences) determine the size of the weighting pool. It is also plausible that pooling sizes vary for different stimulus features. For instance, recent evidence from our lab indicates that the neural representation of multiple speeds of overlapping stimuli in MT aligns with a global pooling scheme (Huang et al., 2023). Understanding these variations could provide deeper insights into the neural representation of multiple visual stimuli.

### Modulation of MT response by object-based (surface-based) attention

Behavioral effects of object-based attention have been demonstrated using transparent motion stimuli (Valdes-Sosa et al., 1998, 2000; Mitchell et al., 2003, 2004). Using similar stimuli, Wannig et al. (2007) showed that the responses of MT neurons can be modulated by object-based attention directed to a moving surface of two overlapping stimuli. Modulation of neuronal responses by object-based attention has also been found in V1 (Roelfsema et al., 1998) and V4 (Fallah et al., 2007). Our study showed new evidence of response modulation by object-based attention in MT. Importantly, our visual stimuli and experimental paradigm allowed us to distinguish the effect of object-based attention from feature-based attention.

Responses of MT neurons can be modulated by feature-based attention to motion direction (Treue and Martinez-Trujillo, 1999; Martinez-Trujillo and Treue, 2004), and binocular disparity (Ruff and Born, 2015). At the neuron population level, attending to the preferred direction of an MT neuron enhances the response to visual stimuli in the RF relative to attending to the null direction, regardless of whether the RF stimulus moves in the preferred or null direction of the neuron (Martinez-Trujillo and Treue, 2004). The same result was found for attending to binocular disparity (Ruff and Born, 2015). These results are consistent with the “feature-similarity gain (FSG) model” of attention (Martinez-Trujillo and Treue, 2004; Maunsell and Treue, 2006). In our study, however, we found that attending to the more preferred disparity of MT neurons did not always enhance MT responses to two moving surfaces. When the surface at a more preferred disparity moved in a less preferred direction and elicited a weaker component response, attending to the more preferred disparity reduced MT response. Conversely, when the surface at a less preferred disparity moved in the preferred direction and elicited a stronger component response, attending to the less preferred disparity enhances MT response (Fig. 10). Therefore, the attention modulation was not due to feature-based attention to binocular disparity. To account for the neural response modulation found in our results, the attention initially directed to a cued binocular disparity of a stationary surface must spread to the motion direction of the attended surface of the overlapping stimuli, although the motion direction was never explicitly cued. Because the motion direction and the cued binocular disparity belong to the same surface, selecting the surface by attention would allow attention to spread to other features of the same object. Whether attention enhanced or reduced the neural response to two surfaces was dependent on whether the attended surface elicited a stronger or weaker component response than the other surface, rather than whether the attended surface was at the preferred or less preferred disparity of the neuron.

Since stereoscopic depth can be considered a spatial dimension in 3D space, may the effect of attention modulation found in our study be due to spatial attention? In response to two 2D stimuli placed at different locations within the RF, the neuronal response in MT (Treue and Maunsell, 1996; Seidemann and Newsome, 1999), V2, and V4 (Moran and Desimone, 1985; Luck et al., 1997; Reynolds et al., 1999) is enhanced (or reduced) when attention is directed to the spatial location where a stimulus component is a preferred (or null) stimulus of the neuron when presented alone. It has been shown that these results can be explained by the biased competition model (Desimone and Duncan, 1995; Reynolds et al., 1999) and the model of divisive normalization (Ghose and Maunsell, 2008; Lee and Maunsell, 2009; Reynolds and Heeger, 2009; Ni et al., 2012). Our finding that attending to a surface eliciting a stronger (or weaker) component response when presented alone enhanced (or reduced) the response to overlapping 3D stimuli is consistent with these previous studies. Our study made two new advancements. First, it extended the finding to 3D stimuli with attention directed to the feature of binocular disparity. Second, previous studies using 2D stimuli did not measure the spatial preferences of the neurons. In contrast, our study directed animals’ attention to the more preferred or less preferred disparity of the neurons and showed that the attention modulation was not dependent on the attended disparity, but dependent on the combined stimulus features of the attended surface as a whole. Regarding whether the attention modulation found in our study was due to spatial attention or object-based attention, we think the distinction may be more semantic than mechanistic. One can think of a surface in our study either as an object or a spatial location in depth. In the previous studies, although attention was directed to a spatial location within the RF, it was also possible that attention selected the visual stimulus as an object at the attended location which allowed spatial attention to spread to other features of the stimulus such as motion direction or orientation. Cavanagh and colleagues (2023) have recently suggested that spatial attention may rely on object-based attention.

### Functional implication of disparity bias on segmentation and figure-ground segregation

It becomes easier to segment two overlapping surfaces when they are moving at different depths than at the same depth (Hibbard and Bradshaw, 1999; Greenwood and Edwards, 2006). The neural representation of two surfaces moving transparently in different directions and at different depths is distributed in the responses of populations of neurons that are selective for motion directions and binocular disparities. Without disparity cues, the visual system can only rely on motion cues to segment the overlapping surfaces (Xiao and Huang, 2015). When two moving surfaces are located at different depths, even when neurons simply average the component responses without showing disparity bias, MT neurons that prefer the disparity of one surface would still carry more information about the motion direction of that surface and therefore aid in segmentation. However, the benefit can be limited when the differences in stimulus features (disparity and direction) are small. The disparity bias that we found would further enhance the neural representation of the motion direction at the surface located at the more preferred disparity of the neurons. This disparity bias, independent of attention modulation, could help further partition the neural representation of multiple stimuli into distributed responses across different neuronal populations and therefore facilitate segmentation. Together, these mechanisms may provide a neural basis for the perceptual benefit of depth cues on segmentation.

Three lines of evidence from our results suggest that the near surface of overlapping stimuli may be better represented than the far surface. First, in monkey G, we found a predominant near bias across near and far neurons. The overall near bias could facilitate the segmentation of the near surface, which is consistent with the better performance of monkey G in discriminating motion directions of the near surface than the far surface of overlapping stimuli. Second, for animals like monkeys B and R, near surface may be better represented than the far surface because more neurons tend to prefer near disparity over far disparity across visual areas of V1, V2, V4 (Prince et al., 2002; Chen et al. 2008; Gonzalez et al., 2010; Samonds et al., 2012; Hinkle et al., 2005; Tanabe et al., 2005), and MT and MST (Maunsell and Van Essen, 1983b; DeAngelis and Uka, 2003; Gonzalez et al., 2001). Since near neurons tend to show near bias and far neurons tend to show far bias, having more near neurons in the population would better represent the near surface of overlapping stimuli. Third, when combining all neurons in our dataset for each of the three animals, we found an initial near bias relative to the average of the component responses (Fig. 9, Table 1). This finding suggests that, at least in the early response, the near surface is better represented than the far surface, making it likely easier for the near surface to be segregated from the far surface, or to be selected by attention. Figural objects in natural scenes are more likely to reside at near disparity than far disparity relative to the ground region (Huang et al., 2019 SFN abstract). A better neural representation of the near surface in the visual cortex during the early neural response may reflect a strategy that the visual system uses to exploit the statistical regularity of natural vision and to better serve the function of figure-ground segregation. The current study only used surfaces at two disparities of -0.1⁰ and 0.1⁰. Future studies using a range of disparities are needed to further test this idea.

## Methods

### Subjects and neural recording

Three adult male rhesus monkeys (Macaca mulatta) participated in the behavioral and neurophysiological experiments. All experimental procedures were conducted in accordance with protocols approved by the Institutional Animal Care and Use Committee of the University of Wisconsin–Madison, adhering to U.S. Department of Agriculture regulations and National Institutes of Health guidelines for the care and use of laboratory animals. The procedures for surgical preparation and electrophysiological recordings were similar to those described in previous studies (Xiao and Huang, 2015).

We implanted a head post and a recording cylinder during sterile surgery, with the animal under isoflurane anesthesia. For electrophysiological recordings from neurons in area MT, we employed a vertical approach, using either single tungsten electrodes (1–3 MΩ, FHC) or 16-channel linear probes (AlphaOmega; Plexon). We identified area MT in each animal using structural MRI, along with its hallmark features: a high proportion of direction-selective neurons, smaller RFs compared to those in the adjacent medial superior temporal cortex (area MST), its specific anatomical location on the posterior bank of the superior temporal sulcus, and the visual topography of the RFs (Gattass and Gross, 1981). We amplified the electrical signals and identified single units using a real-time template-matching system in conjunction with an offline spike sorter (Plexon). Eye position was monitored at a rate of 1000 Hz using a video-based eye tracker (EyeLink, SR Research). In certain recording sessions with monkey B, we tracked both the left and right eye positions, which were subsequently used to calculate horizontal vergence by subtracting the horizontal position of the right eye from that of the left eye.

### Experimental procedures

We used the real-time data acquisition program “Maestro” to control both stimulus presentation and data collection (https://sites.google.com/a/srscicomp.com/maestro). Visual stimuli were displayed on a 25-inch CRT monitor (SONY) positioned at a viewing distance of 63 cm. The monitor had a resolution of 1024 × 768 pixels and a refresh rate of 100 Hz. Stimuli were generated on a Linux workstation using an OpenGL application that communicated with the experimental control computer. Luminance calibration of the CRT monitor was performed using a photometer (Minolta LS-110). Gamma correction was applied and the output linearity was subsequently confirmed.

Once the spike waveforms of a single neuron were isolated, we initially assessed the neuron’s direction selectivity by presenting interleaved trials of a 30° × 27° random-dot patch (RDP), which moved in various directions with 45° increments at a speed of 10°/s. The direction selectivity and preferred direction (PD) were characterized in real time using MATLAB (MathWorks) for data analysis and visualization. Next, we assessed the neuron’s speed tuning by presenting a random dot patch moving at various speeds (1, 2, 4, 8, 16, 32, or 64°/s) in the neuron’s PD. We determined the preferred speed (PS) by fitting the speed-tuning curve with a cubic spline and identifying the speed that elicited the highest firing rate. Following this, we mapped the neuron’s RF by recording responses to a series of 5° × 5° random dot patches moving in the PD at the PS. These patches were systematically positioned across the screen (35° × 25°) in 5° steps without overlap. We then interpolated the raw response map at 0.5° intervals to identify the location that produced the highest firing rate, designating this point as the center of the RF.

We used green and red anaglyph filters (Kodak Wratten filters, #25 and 58) to generate stereoscopic depth on the monitor through horizontal disparity. To create a disparity of +n° (far) or -n° (near), we presented two identical dot patterns, one in green and the other in red, with the center of the red dot pattern offset horizontally by +n° or -n° relative to the green dot pattern. When the animals viewed the stimuli through a green filter over the left eye and a red filter over the right eye, each eye perceived one set of dots, producing a binocular disparity that resulted in depth perception. Crosstalk between the two eyes, as viewed through the filters, was measured at 14.5%, 9.6%, and 9.6% for monkeys G, B, and R, respectively. Crosstalk was calculated as the luminance of red (green) dots viewed through the green (red) filter divided by the luminance of red (green) dots viewed through the red (green) filter, with the listed values representing the average crosstalk for red and green dots. The crosstalk remained relatively high even when using two layers of Kodak Wratten filters. Upon investigating the source of the crosstalk, we found that the CRT monitor emitted a faint white light even when the look-up table (LUT) value for background luminance was set to zero. The background luminance measured through the green and red filters were 1.01 and 0.68 cd/m^2^, respectively. This suggests that the light leaking through the anaglyph filter of one color was likely ambient background light, rather than the dots of the opposite color. Human observers, using the same anaglyph filters, confirmed the perception of clear stereoscopic depth with two overlapping surfaces at different binocular disparities. We evaluated an MT neuron’s selectivity to binocular disparity by recording its responses to a series of random dot patterns moving in the neuron’s PD and at its PD speed (typically between 5° and 20°/s). These RDPs were presented at fifteen different disparities, ranging from -1.5° to +1.5°.

### Visual stimuli

Our primary visual stimuli consisted of two overlapping RDPs moving at different depths within a stationary circular aperture with a 15° diameter. One RDP moved at a binocular disparity of -0.1° (near depth) and the other at 0.1° (far depth) (Fig. 2A-D). The direction separation between the two moving surfaces was set at either 60° or 120°. We refer to each RDP as a “stimulus component,” with one being the near component and the other the far component. The dot density for each RDP was 0.89 dots/deg^2^, and both RDPs moved at 100% coherence. As each dot reached the border of the aperture, it would disappear and reappear on the opposite side, continuing its motion in the same direction until the end of the motion period. We varied the VA direction of the bi-directional stimuli in randomly interleaved trials. We also randomly interleaved trials in which only the near or far component was shown alone.

To maximize the behavioral performance of monkeys B and G on the 12AFC task, we typically centered the visual stimuli on the fixation point. Sometimes, we shifted the stimuli 1 to 2° towards the RFs to better cover the RFs. As a result, the visual stimuli encompassed the MT RFs, though they were not directly centered on them. For monkey R, the bi-directional stimuli and component stimuli were centered on the RFs with a diameter of 15°. Meanwhile, a task-relevant random-dot stimulus with a diameter of 5°, moving in a single direction at the fixation plane, was presented in the opposite visual hemifield, centered 10° to the left of the fixation spot (see behavioral tasks below). The speed of the motion stimuli was typically set at 10°/s, with occasional adjustments to 12°/s. These stimulus speeds drove most MT neurons effectively and facilitated visual segmentation of transparently moving stimuli at these relatively slow speeds.

### Behavioral tasks

Monkeys B and G performed a 12-alternative forced-choice (12AFC) direction discrimination task to report the motion direction of a surface at a cued depth. Each trial began with the animals fixating on a central point, while stationary reference dots at zero disparity were displayed outside the region where the stimuli would later appear. The monkeys had to maintain fixation within a 1° × 1° window. To direct attention, a static circular random dot pattern (RDP) with a 15° diameter appeared for 500 ms at either near (−0.1°) or far (0.1°) disparity. Stationary reference dots at zero disparity filled the rest of the display to enhance depth perception. Following a 300-ms blank screen with the fixation point, a single RDP at the cued disparity or two overlapping RDPs at 0.1° and -0.1° disparities appeared stationary for 500 ms, followed by 600 ms of motion. The VA direction of the motion was one of 12 directions spaced at 30° intervals from 0° to 360°. After the motion period, the stimuli disappeared, and 12 reporting targets, arranged in a 360° circle at 30° intervals with a 10° diameter, appeared for 450 ms for Monkey B and 600 ms for Monkey G. During this time, the monkeys reported the motion direction of the cued component by making a saccade to the corresponding target in the matching direction. They were required to hold fixation within a 1.5° × 1.5° window around the correct target for 350 ms to receive a juice reward.

In the task where attention was directed away from the RFs, single- and bi-directional stimuli at different depths were centered on the RFs of MT neurons, while Monkey R attended to a task-relevant stimulus in the opposite visual hemifield. Monkey R performed a 2AFC fine direction discrimination task, reporting whether the motion of an RDP was clockwise or counter-clockwise relative to an invisible upward vertical. The monkey maintained fixation within a 1° × 1° window until making a saccade to report. The attended stimulus was a 5° diameter RDP, placed 10° to the left of fixation at zero disparity, while RF stimuli were positioned in the right hemifield (recording from the left MT). Each trial began with a fixation point and reference stationary dots at zero disparity. After 200 ms of fixation, both the RF and attended stimuli appeared stationary for 250 ms before moving for 500 ms. The attended stimulus moved at 5°/s, either clockwise or counter-clockwise from vertical by offsets of 10°, 15°, or 20°. The RF stimuli with either a single RDP or overlapping RDPs moved in one of the 12 directions as described above in “Visual stimuli”. All trials were randomly interleaved. After the motion period, stimuli were turned off, and two reporting targets appeared 10° to the left and right of fixation. Monkey R had 400 ms to saccade to the left target if the attended motion was counter-clockwise to the upward vertical, or the right target if clockwise to vertical, to receive a juice reward.

### Data analysis

Neuronal responses were quantified as the firing rate (spikes per second) during the stimulus motion period, averaged across repeated trials. We constructed tuning curves for both unidirectional and bi-directional stimuli, fitting the raw data using cubic splines with 1° interpolation steps. For each VA direction of the overlapping stimuli, we measured the responses to the bi-directional stimuli as well as to the individual unidirectional components. To average the direction tuning curves across neurons, we aligned the spline-fitted tuning curves for bi-directional stimuli by rotating them so that the VA direction of 0° corresponded to each neuron’s PD. We then normalized the responses based on the maximum firing rate to either the near or far component, whichever was greater, and averaged the aligned, normalized tuning curves across all neurons.

We calculated the population-averaged tuning curves by combining responses from the Near C/Far CC and Near CC/Far C conditions (Fig. 3). For visualization, the near component was positioned on the clockwise side and the far component on the counter-clockwise side. To achieve this, we horizontally flipped the spline-fitted, aligned tuning curves for each neuron in the Near CC/Far C condition relative to the vertical axis at VA direction 0° (the neuron’s PD). The flipped tuning curves were then averaged with the tuning curves from the Near C/Far CC condition.

To analyze the time course of response tuning to the bi-directional stimuli, we calculated the tuning curves for each neuron using trial-averaged firing rates within a sliding 50-ms window, stepping by 10 ms. Responses were normalized to the maximum firing rate across all time bins, based on the stronger response to either the near or far component. We then pooled the responses from the Near C/Far CC and Near CC/Far C conditions, as described above, and averaged the normalized responses across neurons.

To evaluate the goodness of fit for a model of response tuning to bi-directional stimuli, we calculated the percentage of variance (*PV*) explained by the model using the following formula:

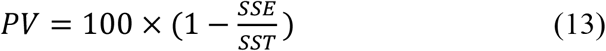

where *SSE* is the sum of squared errors between the model fit and the neural data, and *SST* is the sum of squared differences between the data and the mean of the data.

### Divisive normalization model (DNM)

#### Probability density function of weighting pool size

We fitted each neuron’s response tuning curve to the bi-directional stimuli using an extended divisive normalization model. Each neuron contributed to the fits of eight bi-directional tuning curves presented in the results (2 direction separations of 60° and 90° *×* 2 stimulus configurations of “Near C/Far CC” and “Near CC/Far C” *×* 2 attention conditions of “Attend Near” and “Attend Far”). Equations 11 and 12 were used to fit the bi-directional tuning curves of each neuron. When fitting the response tuning curve with the DNM model, we divided the parameter search space for the parameter *v* (the standard deviation of the Gaussian function ***G***) into steps that resulted in equal steps of the area under the truncated Gaussian function ***G*** (Eq. 10) (within -1.5° to 1.5° disparities) (Eq. 10). We did so because the area under ***G*** better represents the weighting pool size than *v*, given the nonlinear relationship between *v* and the area. For each *v* value in the search space (ranging from 0.01 to 40 in 20 steps), we used MATLAB function “fmincon” to minimize the sum of squared errors (SSE) between the model fit and the neural data (Eqs. 11 and 12). The steps of *v* were equilvalent to even steps of normalized area under ***G*** from 0 to 1 in a step of 0.05.

We calculated the probability distribution of the weighting pool size that gave rise to the best fit of the data (measured by SSE) using the DNM. To do so, we first calculated the probability distribution of *v*, then converted it to the probability distribution of the area under the Gaussian function ***G***. For each neuron, we first calculated the SSE of the model fit as a function of *v*. Instead of selecting a single *v* value with the lowest SSE, we considered all values of *v* that produced SSE within the lowest 5%, as multiple *v* values could yield similar fits. Given the inherent noise in neuronal responses, it was important to account for all *v* values that provided a good fit. For each tuning curve, we normalized SSE (*v*) by the maximum SSE and defined a function *f(v)* such that *f(v)* = 1 when the normalized SSE at those *v* values were ≤ 0.05, and *f(v)* = 0 at other *v* values. We then normalized *f(v)* by its total area to obtain 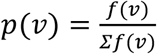. The probability density function *pdf(v)* was computed by averaging *p(v)* across all tuning curves from each neuron and further averaged across near or far neurons. Finally, we converted *pdf(v)* to the probability density function based on the area under the Gaussian function ***G*** (Fig. 11B and C).

To evaluate whether *pdf(v) differed* between the monkeys, we used bootstrapping where we randomly sampled the *pdf(v)* from a neuronal set of either the near neurons or far neurons 1000 times with replacement and calculated the 95% confidence intervals for each *v* value.

#### Estimate model parameters for the effect of attention

We derived the values of attention parameters *β*_*An*_ and *β*_*Af*_ used in Equations 11 and 12 using the following approach. We first fitted the response tuning to the bi-directional stimuli with the LWS model:

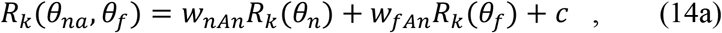

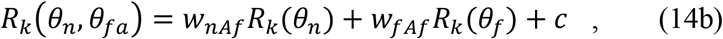

and empirically determined the parameters *w*_*nAn*_, *w*_*fAn*_, *w*_*nAf*_, and *w*_*fAf*_, which are the weights for the near and far components in the two attention conditions. The first subscripts *n* and *f*refer to the near and far components, respectively. The subsequent subscripts *An* and *Af* denote attending near and attending far components, respectively. The parameter *c* is positive.

We assumed that attention to a stimulus component of the bi-directional stimuli multiplicatively scaled the response weight of the attended component (i.e. scaled by factor *β*_*An*_ or *β*_*Af*_), but did not change the weight of the unattended component (i.e. scaled by a factor of 1).

So *w*_*nAn*_, *w*_*fAn*_, *w*_*nAf*_, and *w*_*fAf*_ can be expressed as the following. When attention was directed to the near component:

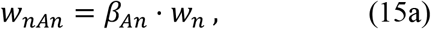

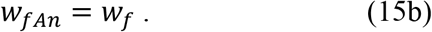

When attention was directed to the far component:

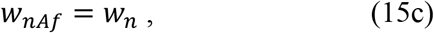

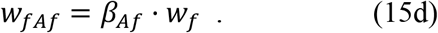

*w*_*n*_ and *w*_*f*_ are the response weights for the near and far components, respectively, when attention is not involved. Therefore, by combining Equations 15a and 15c, we can determine *β*_*An*_ *(Eq. 16a)*.

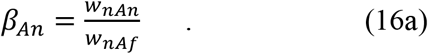

The attention factor *β*_*An*_ is essentially a ratio between the response weights for the near component under the attending near and attending far conditions. Since *w*_*nAn*_ and *w*_*nAf*_ can be estimated from the data, *β*_*An*_ can be determined accordingly. Similarly, by combining Equations 15d and 15b, we can determine *β*_*Af*_ *(Eq. 16b)*.

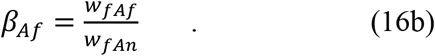

## Supplementary Materials

## Methods and Results

We performed additional analysis to examine the timecourse of disparity bias and its relationship to neuronal responses. To track the neuronal response to bi-directional stimuli (*R*_*12*_), we averaged the responses from the ‘Attend Near’ and ‘Attend Far’ conditions for monkeys B and G, and used the ‘Attend Away’ response for monkey R. The firing rate was calculated using a 50-ms sliding window, advancing in 10-ms steps. Within each time window centered on time point *t*, we integrated the neuronal response within a 30° direction window centered on the VA direction at the “Near PD” and “Far PD” regions of the neural responses from monkeys B, G, and R, as shown in the colormaps in Figures 6, 7, 8. The integrated responses, denoted as *R(NearPD, t)* and *R(FarPD, t)*, were averaged to estimate the timecourse of the overall neuronal response (blue curves in Supplementary Figures 1-3).

To capture the timecourse of disparity bias, we calculated the difference between *R(NearPD, t)* and *R(FarPD, t)* (purple curves in Supplementary Figures 1, 2, and 3). *R* was derived either from the bi-directional response *R*_*12*_ (thick purple lines) or from the difference between *R*_*12*_ and the average of the component responses *R*_*avg*_, (i.e. *R*_*12*_ - *R*_*avg*_) (thin purple lines).

For monkey B, relative to the neuronal response, the near bias of the near neurons (SFig. 1A, B) appeared sooner than the far bias of the far neurons (SFig. 1C, D). The disparity bias also appeared sooner at 120° DS (DS) (SFig. 1B, D) than at 60° DS (SFig. 1A, C). At 120° DS, the near bias of the near neurons almost had no delay (SFig. 1B). Interestingly, at 60° DS, the far neurons initially showed a weak near bias before changing to the far bias (the thin purple curve in SFig. 1C).

For monkey G, the neuronal response was more delayed relative to monkey B. The disparity bias of monkey G was also substantially more delayed (SFig. 2) than monkey B (SFig. 1). Both near neurons and far neurons of monkey G showed near bias. Similar to monkey B, the near neurons of monkey G (SFig. 2A, B) showed a sooner disparity bias than the far neurons (SFig. 2C, D).

Similarly, the near neurons of monkey R showed a sooner near bias (SFig. 3A, B) than the far neurons showing a far bias (SFig. 3C, D). The near bias of the near neurons also rose faster than the far bias of the far neurons (comparing SFig. 3A with 3C, and 3B with 3D). As found in monkey B, the far neurons from monkey R initially showed a near bias, before gradually changing to the far bias (thin purple curves in SFig. 3C, 3D).

**SFigure 1.**
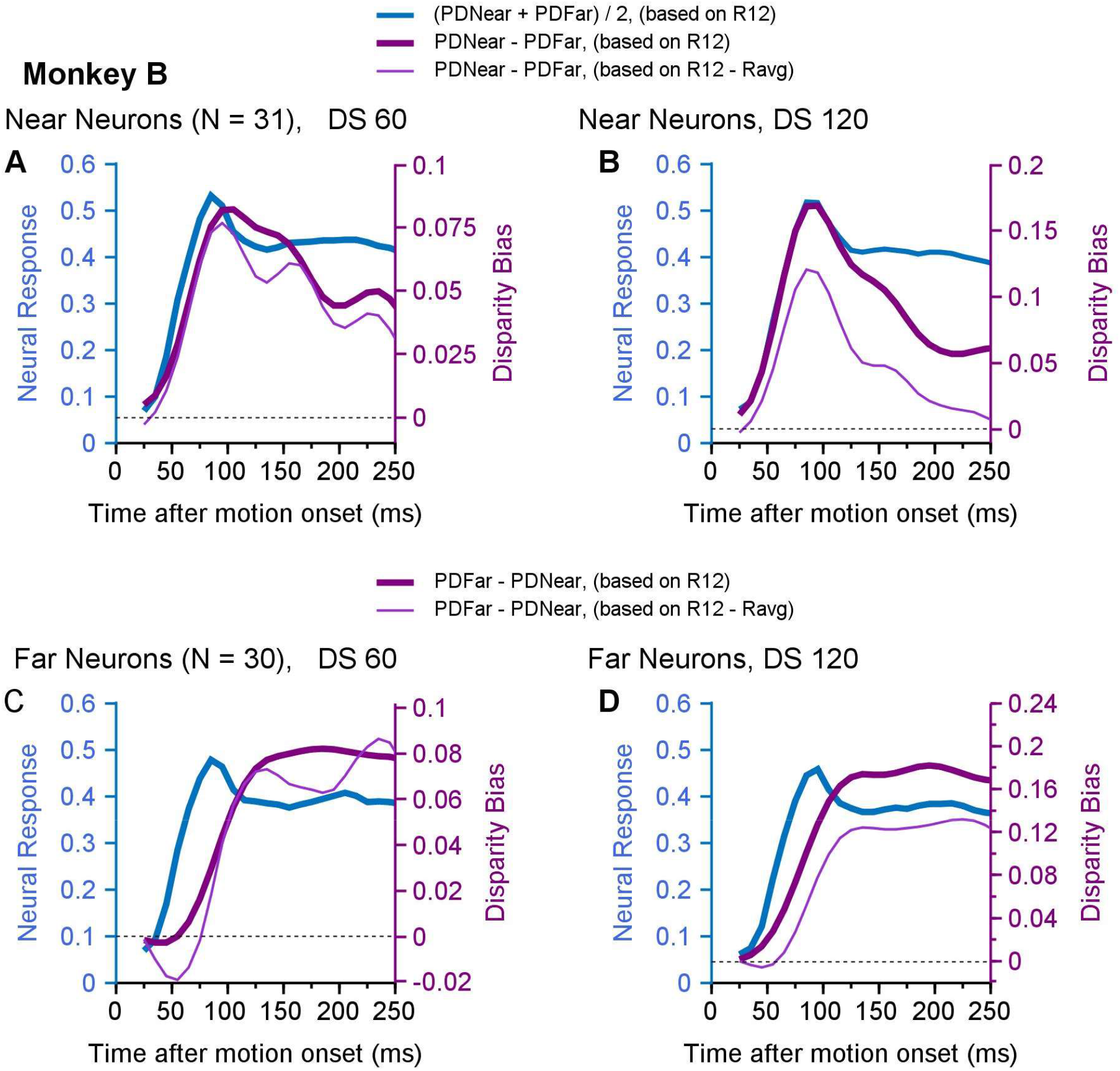
Comparison of the timecourses of neuronal responses and disparity bias in monkey B. The blue curves show the population averaged neuronal response to bi-directional stimuli (*R12*) measured by averaging *R(NearPD)* and *R(FarPD)*. The thick puple curves indicate the disparity bias by comparing *R(NearPD)* and *R(FarPD)*. The thin puple curves indicate the disparity bias based on *R12* – *Ravg*. The magnitude of the disparity bias is shown by the Y-axis on the right side of each plot, marked by purple color. **A, B**. Responses from near neurons. Disparity bias was calculated by *R(NearPD)* - *R(FarPD)*. **C, D**. Responses from far neurons. Disparity bias was calculated by *R(FarPD)* - *R(NearPD)*. Negative disparity bias below the dotted lines in C, D, indicates near bias. **A, C**. 60° direction separation. **B, D**. 120° direction separation.

**SFigure 2.**
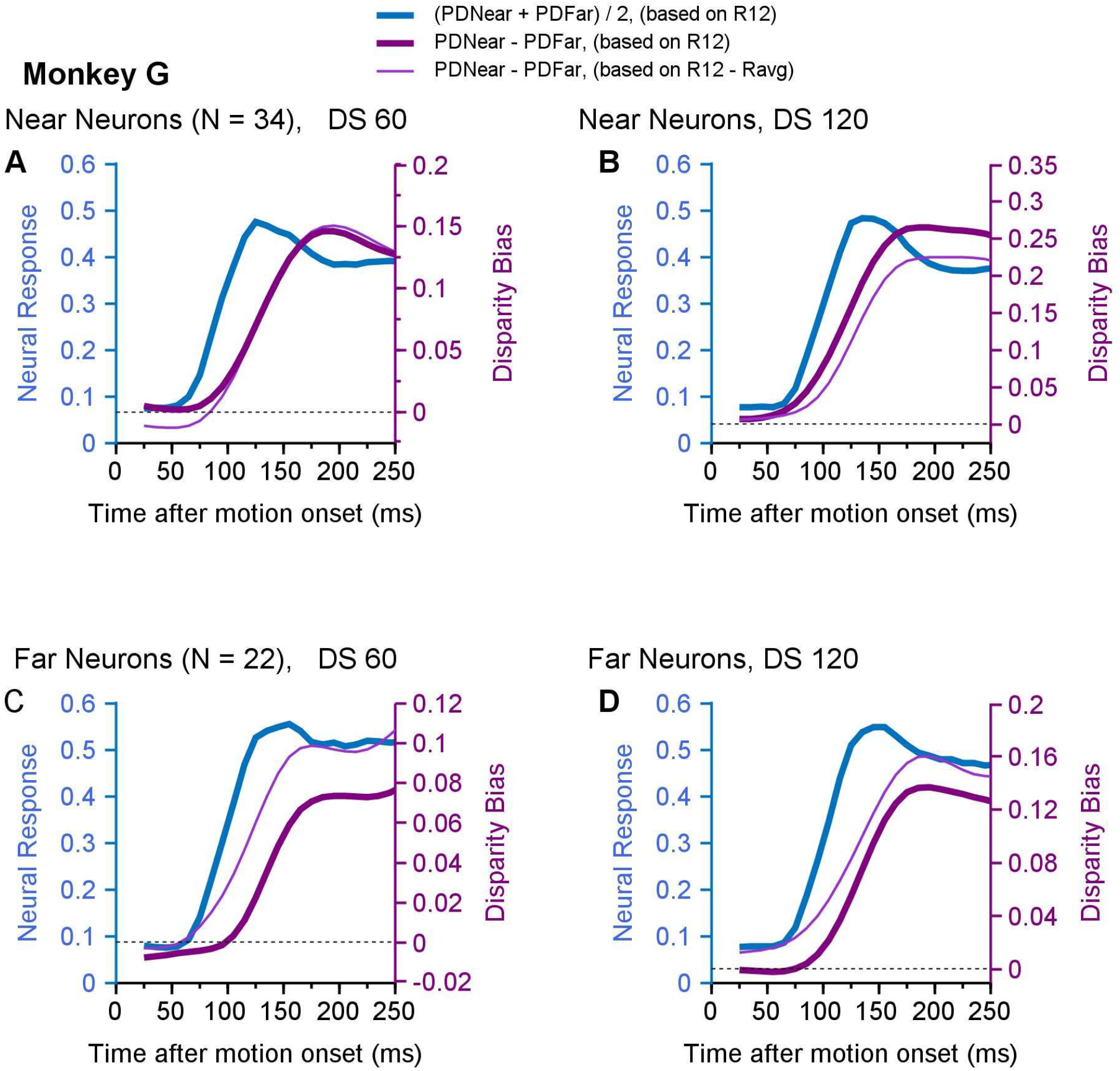
Comparison of the timecourses of neuronal responses and disparity bias in monkey G. The convention is the same as Sfigure 1. **A, B**. Responses from near neurons. **C, D**. Responses from far neurons. Because the far neurons of monkey G also showed near bias, the disparity bias in C, D was calculated by *R(NearPD)* - *R(FarPD)*. **A, C**. 60° direction separation. **B, D**. 120° direction separation.

**SFigure 3.**
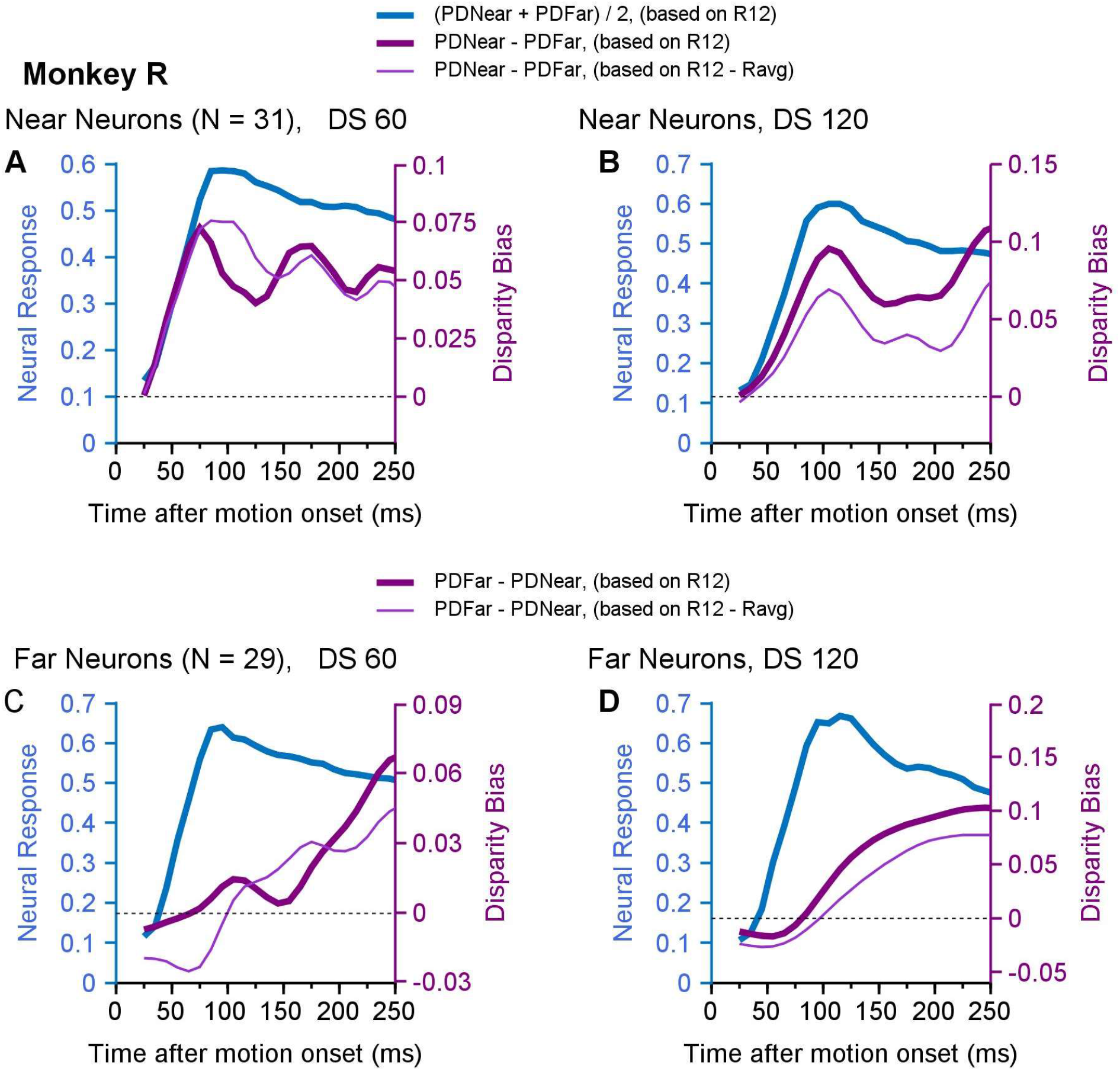
Comparison of the timecourses of neuronal responses and disparity bias in monkey R. The convention is the same as Sfigure 1. **A, B**. Responses from near neurons. Disparity bias was calculated by *R(NearPD)* - *R(FarPD)*. **C, D**. Responses from far neurons. Disparity bias was calculated by *R(FarPD)* - *R(NearPD)*. Negative disparity bias below the dotted lines in C, D, indicates near bias. **A, C**. 60° direction separation. **B, D**. 120° direction separation.

## Acknowledgment

We thank Dr. Greg DeAngelis for sharing the data set of the disparity tuning curves of a large number of MT neurons from his lab to be used in our model, Bryce Arseneau for animal training, and Drs. Jennifer Coonen and Kevin Brunner at the Wisconsin National Primate Research Center for excellent veterinary care and surgical assistance.

